# Multipair phase-modulated temporal interference electrical stimulation combined with fMRI

**DOI:** 10.1101/2023.12.21.571679

**Authors:** Iurii Savvateev, Florian Missey, Valeriia Beliaeva, Sofia Peressotti, Marija Markicevic, Diana Kindler, Fabrice Chaudun, Giulia Casarotto, Camilla Bellone, Christian Lüscher, Daniel Razansky, Viktor Jirsa, Adam Williamson, Rafael Polania, Valerio Zerbi

## Abstract

Temporal Interference stimulation (TIS) is a promising non-invasive brain stimulation technique exploiting frequency-shifted kHz fields to modulate oscillatory neural activity without invasive procedures. While human studies suggest TIS efficacy in targeting relatively deep brain structures, recent computational modeling and animal studies indicate potential off-target stimulations via the application of standard TIS protocols. Here, we computationally optimized TIS targeting for the prefrontal cortex (PFC) in mice. Combining *in vivo* electrophysiological recordings, intracellular calcium dynamics with fiber photometry, and functional MRI, we confirmed TIS-induced amplitude modulation, neuronal entrainment and hemodynamic responses in the PFC, while also identifying off-target modulations. To mitigate off-target effects, we propose a novel configuration with three electrode pairs, one of which is actively phase-shifted by 180 degrees. This cancelling field significantly enhanced TIS focality, reducing off-target effects without compromising the TIS efficacy in the target area. This work effectively addresses one of TIS most critical shortcomings and opens new avenues for research and clinical applications.

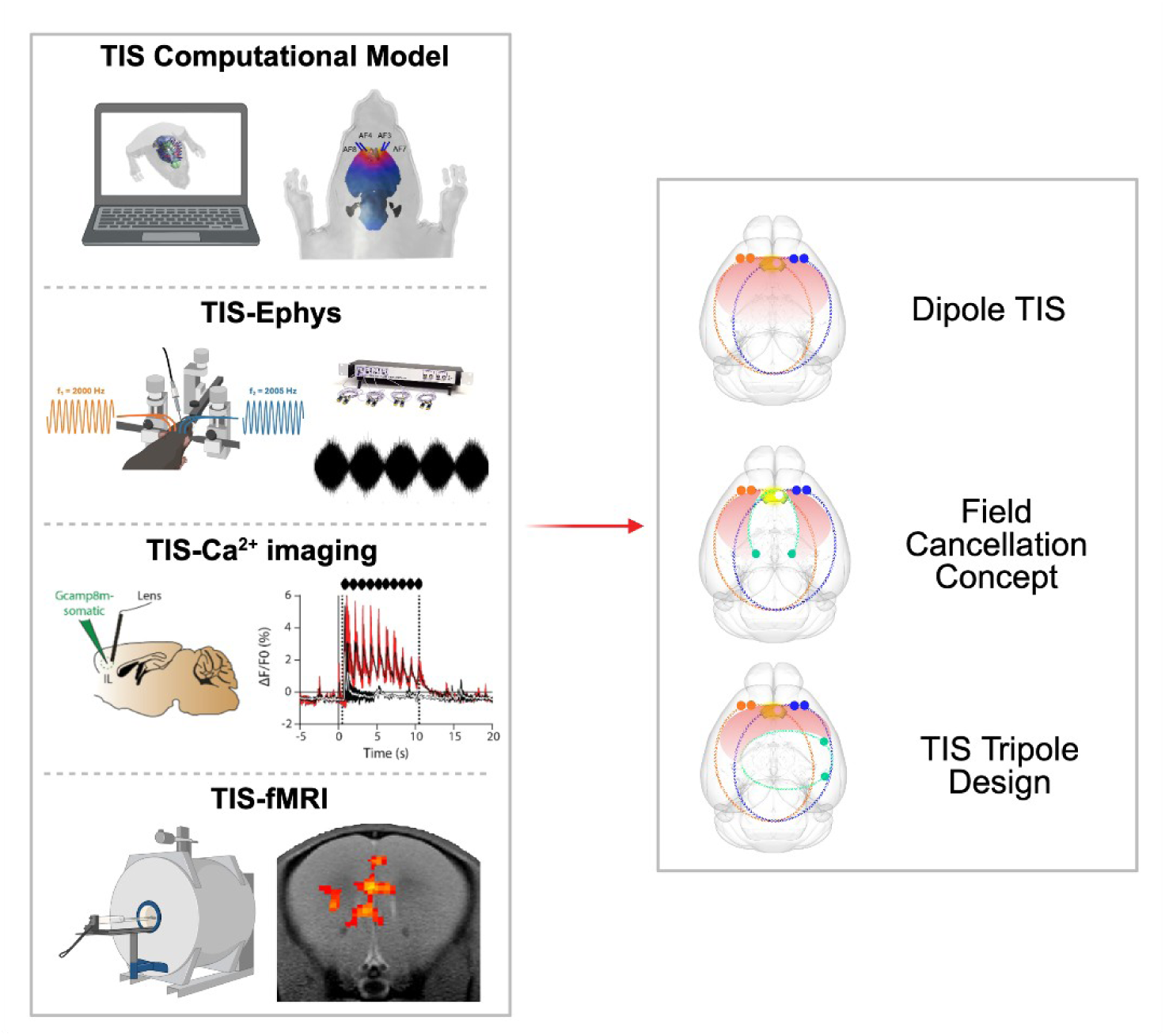

## Introduction

Our understanding of the interplay between brain and behavior has been significantly advanced by non-invasive brain stimulation approaches. Transcranial magnetic stimulation (TMS) and transcranial direct stimulation (tDCS) or alternating current stimulation (tACS) have gained increasing popularity in recent years due to their potential in treating psychiatric disorders such as depression^1–5^ and obsessive compulsive disorder (OCD)^6^. However, non-invasive transcranial stimulation methodologies are primarily employed to affect cortical structures, as opposed to deep brain regions, because of their limited depth penetration^7^.

Invasive brain stimulation techniques, such as deep brain stimulation (DBS), entail electrode implantation into the brain ^4^. Kindled by the therapeutic usage of DBS at the second half of 20th century^8^, DBS use has currently expanded to the treatment of various psychiatric and neurodegenerative disorders such as OCD, depression, addiction, Tourette and Parkinson Disease^9–11^. Invasive brain stimulation techniques are more precise and effective in targeting specific brain regions and circuits, but they carry a higher risk of complications, infections, bleeding, and damage to the surrounding brain tissue^12^. Thus, the trade-off between invasiveness and spatial precision has hindered clinical adoption of both invasive^4,5^ and non-invasive^1–3,6^ methodologies.

In recent years, a novel non-invasive brain stimulation protocol termed temporal interference stimulation (TIS) was developed, allowing the stimulation of subcortical structures without entraining the activity of the overlying cortex^13–15^. TIS uses scalp electrodes to deliver two high-frequency electric fields (f1 and f2 > 1 kHz). As these fields propagate through the brain, they interact to form an interference pattern: in some regions, the fields align in phase and their amplitudes sum (constructive interference), while in others, they oppose and partially cancel out (destructive interference). This spatially-dependent interference produces an amplitude-modulated envelope oscillating at the frequency difference Δf = f2 - f1, typically in the physiologically relevant Hz range. It is this low-frequency envelope that is thought to influence neuronal membrane potentials, as opposed to the high-frequency carriers, which are generally considered too fast to modulate neural activity^13^ (but see ^16,17^ for alternative views on the underlying mechanism).

Recently, a number of *in silico* ^16,18–21^ and *in vivo* studies in animals ^22–24^ and humans have demonstrated the TIS effectiveness in functionally stimulating deep brain areas and non-brain structures, such as the hippocampus ^15,25–27^ the superior colliculus ^26,28^, the striatum ^29,30^ the sciatic nerve ^17,31^ and the bladder ^32^. Despite the success of these applications in a range of diseases and the ongoing clinical trials (NIH: NCT05805215, NCT03747601, NCT04761471) substantial doubts about the specificity and off-target effects of TIS remain. For example, an *in silico* study analyzing the biophysics of TIS predicted the occurrence of conduction blocks and unintended stimulation in off-target regions^16^. These findings were later corroborated by *in vivo* experiments, where increasing TIS amplitude by more than 1.5 mA to target the hippocampal CA3 region induced off-target activation in the motor cortex, resulting in aberrant motor behaviors and convulsions in mice^25^. Additionally, Acerbo et al. recorded local field potentials (LFPs) during TIS of the anterior hippocampus in human cadavers and detected envelope signals not only in the targeted area, but also in the central hippocampus and overlying cortex^26^.

In light of this evidence, multiple strategies have emerged to increase the focality of TIS, such as modifying the carrier frequency, reducing the envelope amplitude or introducing multiple supplementary fields (multipair TIS)^18,28^. The multipair mTIS strategy represents a promising approach, although recent evidence is restricted to computational investigations^18,33,34^, peripheral nerve studies^17,35^, and limited empirical validation in brain structures^36,37^.

This study addresses two primary objectives. First, we aimed to validate the core principles of TIS both *in silico* and *in vivo*, providing independent evidence of its efficacy in modulating neural activity non-invasively through electrode configuration and stimulation parameters. As part of this validation, we characterized the dependence of neural responses on stimulation amplitude and carrier frequency and demonstrated field steering effects consistent with previous descriptions of TIS. These experiments established a robust framework for further analysis and ensured that the observed effects were attributable to TIS-specific mechanisms.

Building on this foundation, we investigated the spatial distribution of TIS-induced electric fields across the brain using electrophysiology, functional MRI, and fiber photometry. Notably, we identified off-target field propagation, offering new insights into the spatial constraints of conventional TIS approaches and highlighting the need for improved focality.

To address this limitation, we finally developed a novel multipolar TIS configuration designed to reduce off-target activation through active-phase cancellation. Unlike traditional multi-pair TI strategies, our approach introduces a third, anti-phase interference field that selectively dampens kHz-frequency components outside the target region. This targeted suppression enhances spatial precision and represents a refined strategy for improving the focality of TI-based neuromodulation.

## Results

### Effective neuromodulation response to TIS is validated in silico and in vivo

We first set out to effectively implement TIS, to confirm its reproducibility and characterize its neuromodulation capabilities. As a target, we selected the mouse medial prefrontal cortex (mPFC), a region critically involved in high-level cognition and reward-seeking strategies. This area was chosen for its high translational relevance: abnormal mPFC activity is implicated in various neuropsychiatric conditions, including major depressive disorder, and is a frequent target in clinical DBS protocols ^38^. Demonstrating the ability to non-invasively modulate mPFC activity using TIS would therefore represent a meaningful advance toward clinical translation. We began by implementing an optimal electrode configuration to target the mPFC. A computational modeling pipeline was devised using Sim4Life (Zurich Med Tech AG, Switzerland). The coordinates of our target ROI were based on Allen Brain Common Coordinate Framework (CCFv3) and centered in the infralimbic cortex (IL), a subregion of the mPFC^39^. Overall, 38 electrodes (19 pairs) were virtually placed on the scalp according to a high-density EEG system^40^(**Fig 1A**). TIS fields were simulated, and their efficacy in targeting the IL was assessed for all the electrode configurations. Efficacy was evaluated via two parameters: activation threshold of the IL, and stimulation focality. The threshold corresponded to the median TIS field within the IL, and the focality was calculated as the ratio between the volume above threshold in the IL, and in the whole brain. Finally, a database containing threshold and focality values for each simulated TIS field was created. We identified two optimal electrode pairs (at EEG positions AF3/AF7 and AF4/AF8), which were generating the TIS fields with the highest focality (**Fig 1B**).

**Fig. 1.**
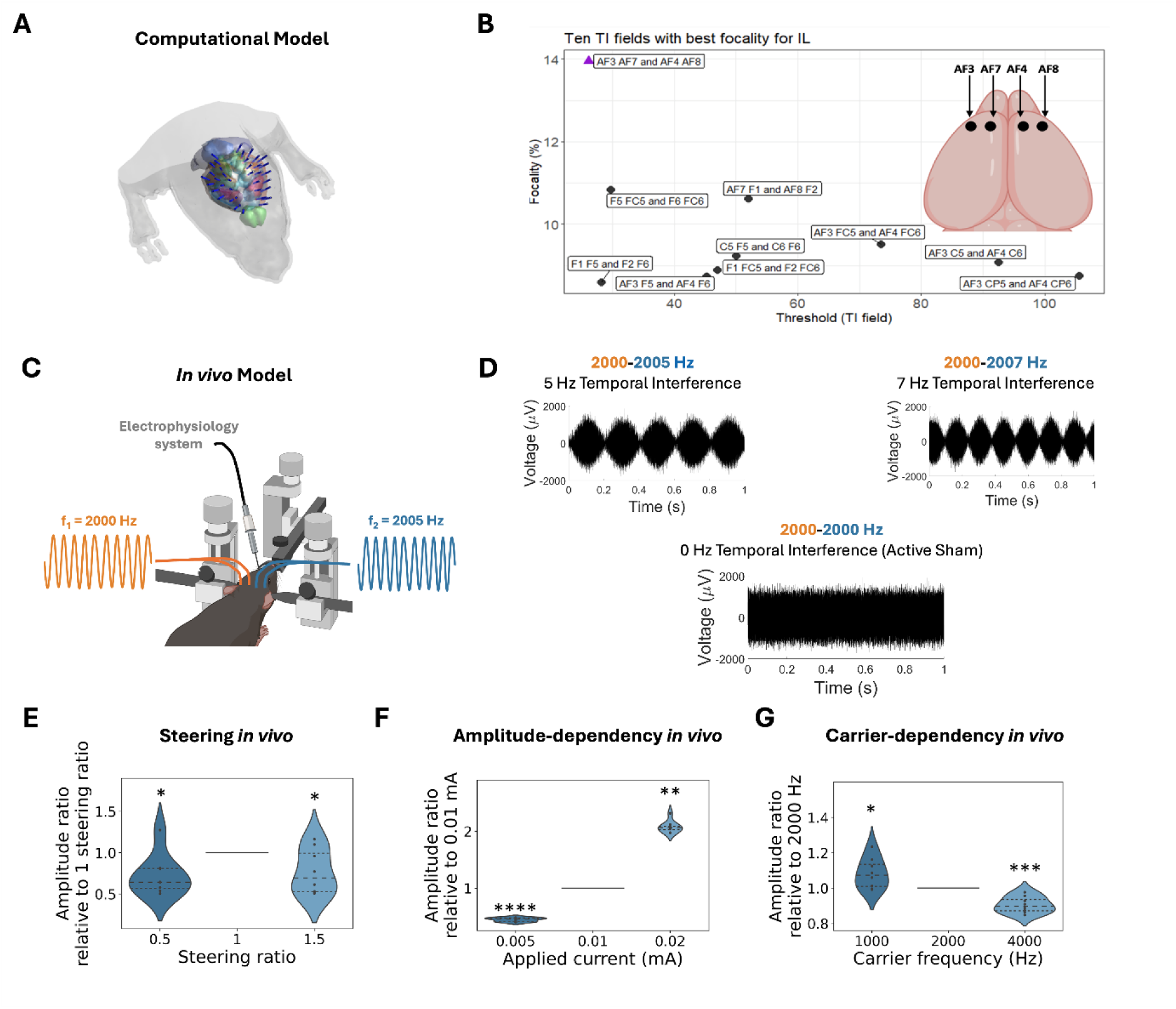
Characterization of the TIS amplitude modulation effect. **(A)** Highly segmented mouse head model showing potential positions for scalp-mounted stimulation electrodes **(B)** Ten electrode configurations with the most optimal focality values for stimulation of the IL region from Sim4life computational modelling. The TIS field generated with AF3, AF7 and AF4, AF8 electrode configuration showed the highest focality. **(C)** Experimental setup for the TIS stimulation (see Methods for details) **(D)** Raw data examples of the envelope modulation (5 and 7 Hz) and active sham recorded in IL *in vivo*. **(E)** LFP measurements of the steering of TIS amplitude outside of the IL region using different current ratios (*p<0.05, n=8, to compare 0.5:1 and 1.5:1 amplitude ratio to 1 one sample Wilcoxon sing-rank test and one sample t-test respectively). **(F)** LFP measurements of scaling of the TIS amplitude in the IL region by changing currents (****p<0.0001, **p<0.01, n=8, to compare amplitude ratios at 0.005mA and 0.02mA to 1 one sample t-test and Wilcoxon sing-rank test respectively). **(G):** LFP measurements of the dependence of TIS amplitude in the IL region from the carrier frequency (*p<0.05, ***p<0.001, n=8, to compare amplitude ratios at 1000Hz and 4000Hz to 1, one sample t-tests).

We then measured TIS-induced activity in the IL region using the electrode configuration selected, to validate the results experimentally. We expected that TIS would elicit neural activity in the IL via amplitude modulation of the two interfering fields (f_1_ and f_2_). The induction of TIS was measured in anesthetized mice *in vivo* and in sacrificed mice *ex vivo* (**Fig 1C, Fig S1**). We applied two 5 µA alternating current (AC) waves at 2000 and 2005 Hz, to generate a 5 Hz amplitude modulation (**Fig 1D**), and recorded the local field potential (LFP) in the IL region. We used an active sham, where both electrode pairs delivered 2000 Hz AC, as a control against the TIS amplitude modulation effects. We showed *in vivo* and *ex vivo* that an amplitude modulation envelope can be generated by the frequency-shifted TIS fields in the IL region, and that the kHz stimulation alone (i.e., the active sham) does not induce any amplitude modulation (**Fig 1D, Fig S1A**), similarly to Grossman et al ^13^.

We then aimed to assess the impact of different current ratios and parameters, to further validate the TIS concept. Changing current ratios of the TIS fields to shift the AM envelope is a method called steering, by which the stimulation target can be adjusted on-demand without repositioning the electrodes. We expected a reduction in TIS stimulation in the IL region when using unequal current ratios, as the electrode configuration had been optimized to target the IL region with equal current amplitudes (ratio: 1:1). Our experimental findings validated this assumption by demonstrating a decrease in TIS amplitude for both the 0.5:1 and 1.5:1 steering ratios, *in vivo* and *ex vivo* (**Fig 1E** and **Fig S1B**). Lastly, we explored the relationship of TIS-induced activation with envelope amplitude and kHz carrier frequency. A range of three amplitudes (AM amplitude: 0.005, 0.01, 0.02 mA) and carrier frequencies (1000, 2000, 4000 Hz) were delivered, and the response was again recorded with LFP in the IL region. We reported a direct proportionality between TIS-induced activation of the IL region and envelope amplitude of the TIS fields (**Fig 1F** and **Fig S1D**). Conversely, TIS activation displayed an inverse relationship with the kHz carrier frequency (**Fig 1G** and **Fig S1C**), exhibiting an IL activation decrease upon increasing the frequency of the carrier, which is consistent with earlier works on mice ^13^ and monkeys ^24^. However, this inverse correlation between higher kHz frequencies and stimulation amplitude could also be explained by hardware limitations under high-impedance conditions (∼10 kΩ), consistent with prior reports of ∼10% output drop between 1–4 kHz ^41^.

### TIS induces neuronal entrainment in the IL across frequencies and amplitudes

After validating the presence of TIS amplitude modulation in the IL region via LFP recordings, it is essential to verify TIS-driven neuronal entrainment. We monitored intracellular calcium dynamics in the IL region with fiber photometry (**Fig 2A**). We delivered 10-s TIS or active sham sessions (**Fig 2B**) at different frequencies (Δf: 0, 0.5, 1, 2, 5, 7 Hz as in **Fig 2C**) and amplitudes (current per pair: 0.125, 0.250. 0.375, 0.500, 0.750 mA). By step-wise increases in 1 Hz TIS current amplitude, we identified the regime at which TIS entrains the IL region for the entire 10 s session (0.750 mA per pair), as observed by the frequency of the fluorescence sensor signal matching the frequency of TIS (in red, **Fig 2D**). Active sham exhibited no periodic changes (in black, **Fig 2D**). Using 0.750 mA current amplitude, we then varied the frequency of the TIS amplitude modulation (**Fig 2C**) to assess whether the detected neuronal entrainment is frequency dependent. At frequencies 0.5, 1, 2 and 5 Hz we detected a frequency match between TIS and the photometry signal, suggesting successful neural entrainment in the IL region across frequencies (**Fig 2E**). At 7Hz, we could observe an initial TIS effect but no continuous entrainment. We observed an initial calcium peak at the start of stimulation in the active sham and TIS conditions, which is a known onset response to kHz frequency fields ^42^. Nevertheless, by detecting calcium activity with fibre photometry, we confirmed TIS-induced neural entrainment, and identified a minimum threshold for the current amplitudes at which we can ensure entrainment at a physiological frequency range.

**Fig 2.**
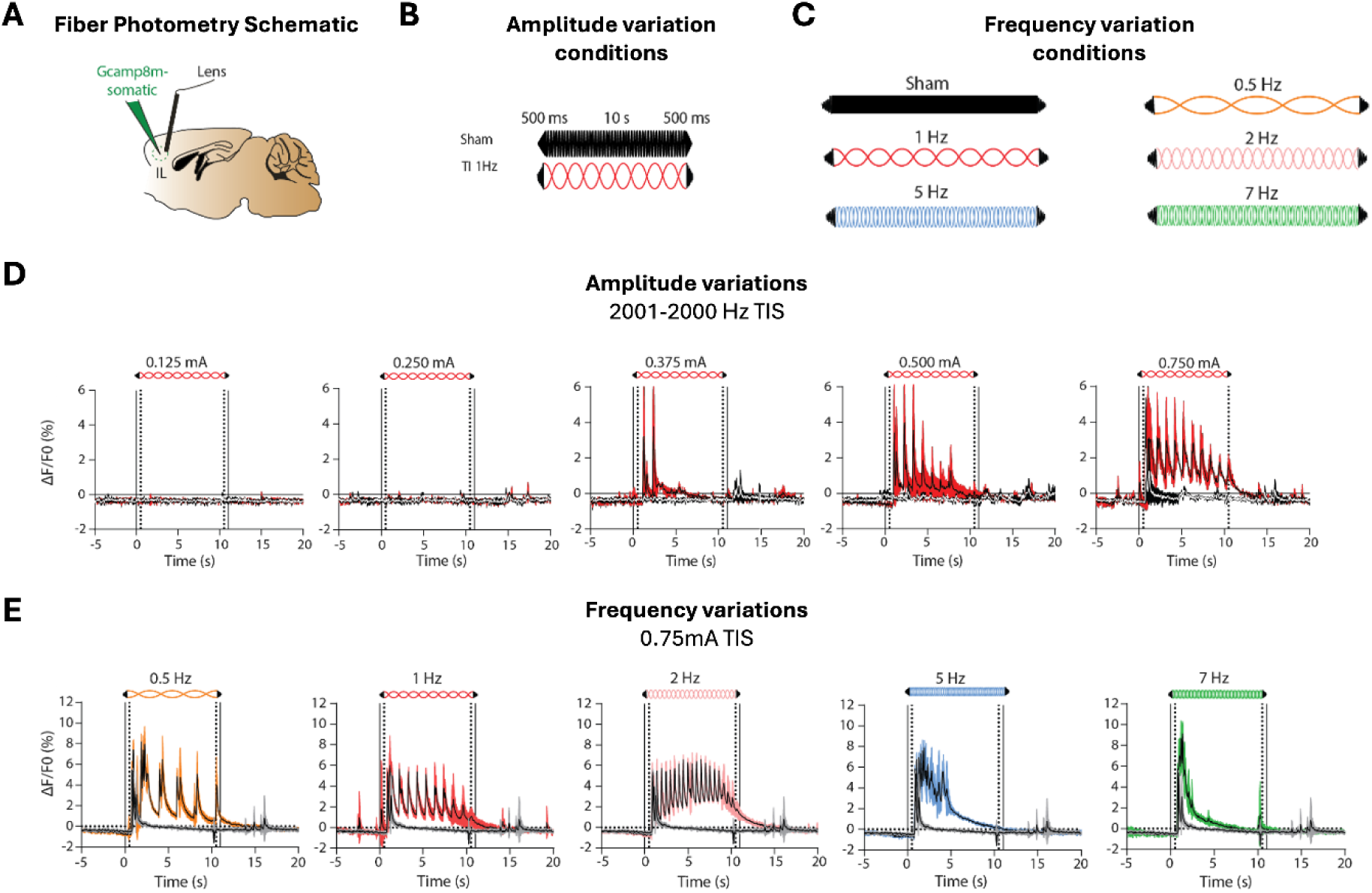
Neuronal entrainment by TIS. **(A)** Schematic representation of fibre photometry design targeting the IL region. **(B-C)** Graphical representations and abbreviations used for intensity- and frequency-dose response experiments. **(D)** Fluorescence signal in the IL region at 1 Hz TIS (2001-2000Hz) at different amplitudes (*n*=2, mean+/-SEM). **(E)** Fluorescence signal in the IL region for different TIS frequencies at the fixed amplitude: 750µA per pair (*n*=1, 5 trials, Mean +/-SD).

### TIS enables spatially selective neuromodulation despite residual off-target signals

After confirming neuronal entrainment from the TIS amplitude modulation, we characterized TIS spatial selectivity (i.e., focality) to deepen our understanding on potential TIS-related off-target activation. We performed electrophysiological recordings in a range of brain regions, and we measured TIS response at the whole-brain level with fMRI. Given the ongoing debate regarding the fundamental mechanisms of TI ^16,17,43^, this TIS-fMRI paradigm in animals is of critical importance, as it constitutes previously uncovered, unbiased evidence of whole-brain mapping during TIS activation.

We first selected four brain regions to record off-target activity, namely the proximal primary motor cortex (MOp) and distal hippocampus (HP) areas, both ipsilateral and contralateral to the IL region. LFPs in the targeted IL area and off-target (MOp(ipsi)/ MOp(contra), HP(ipsi)/HP(contra)) regions were recorded upon 5µA TIS. As predicted by the computational model, the TIS response amplitude in the IL region was higher compared to all other off-target areas, and it decreased along the AP axis (**Fig 3A**). However, the recorded TIS amplitude in MOp and HP were statistically significantly higher than active sham (p=0.0078, paired wilcoxon signed rank test for TIS-active sham in MOp(ipsi), MOp(contra), HP(ipsi), HP(contra) regions, adaptive two-stage Benjamini-Krieger-Yekutieli post-hoc correction, n=8) (**Fig 3B**). To ensure that the TIS amplitude in MOp and HP does not result from the propagation of TIS from the IL region through brain tissue conduction (e.g. axonal projections), we measured phases of TIS oscillations at MOp and HP relative to the IL region (**Table S1**). The detected phase shifts were clustered at 0°, 180°, 360°, suggesting that the corresponding TIS in these areas happened due to passive induction, as opposed to active signal propagation ^44,45^. Thus, while we were able to elicit a significant electrophysiological activation of the IL region using optimized electrode placements and parameters, TIS fields were diminished but not abolished in distant and off-target brain areas.

**Fig. 3.**
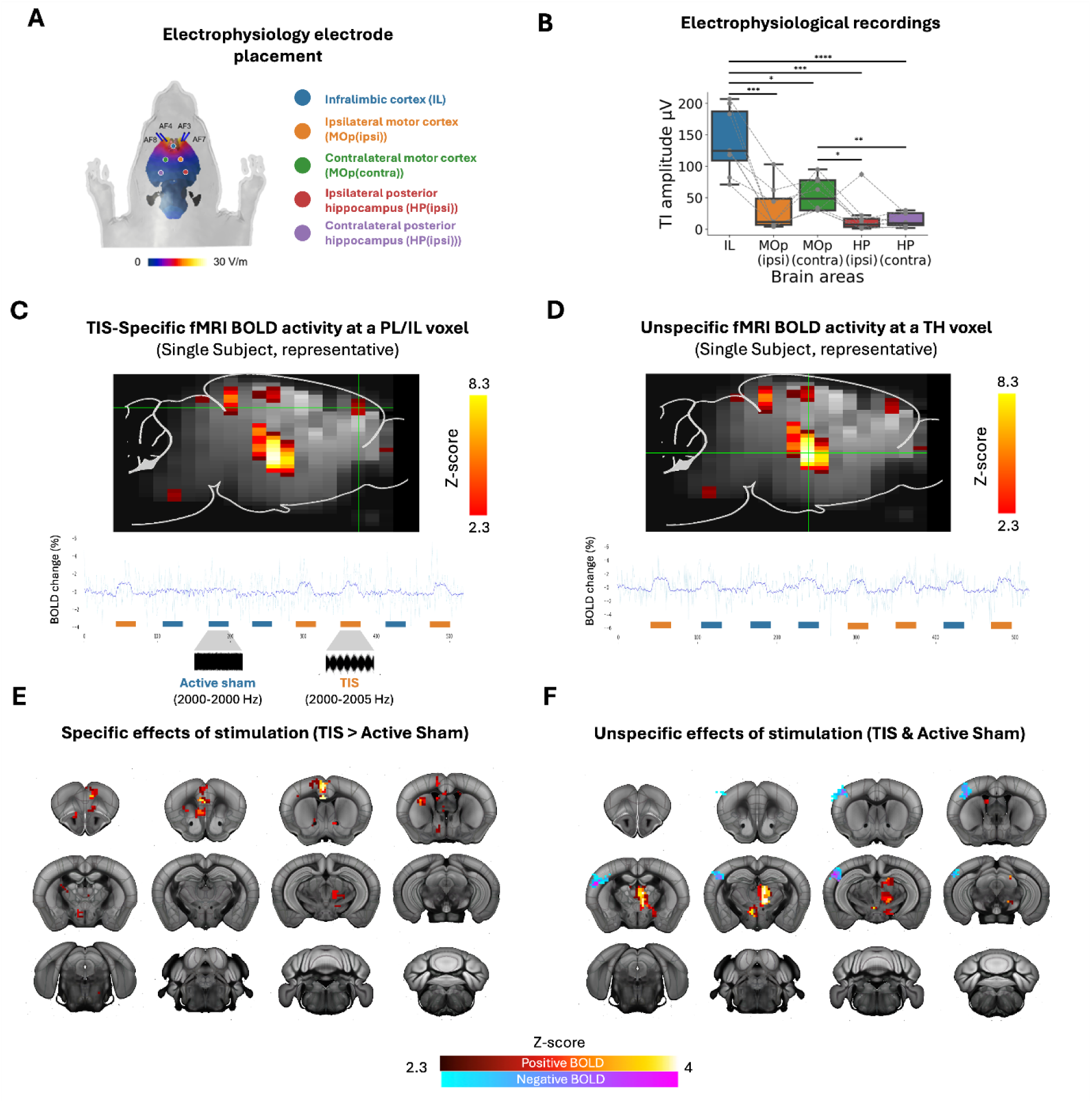
Characterization of TIS focality at the local and whole-brain level. **(A)** schematic of the experimental methodologies used to assess TIS focality. **(B)** Location of stimulation (dark blue) and all recording (light blue, yellow, green, purple, red) electrodes superimposed on the model prediction of TIS field (heatmap)i. The IL target is highlighted in light pink on the heatmap. **(C)** Average TIS amplitudes at the recorded brain areas with the color matching from (B), (*p<0.05, **p<0.01, ***p<0.001, ****p<0.0001, n=8, Friedman test followed by Quade post-hoc test, FDR correction with adaptive two-stage Benjamini-Krieger-Yekutieli method). **(D-E)** Representative Echo Planar Images (EPI) and BOLD timeseries for single animals, from different voxels of a TIS & active sham correlation map, where TIS-specific activation is shown in the target area, loxel PL/IL (D) and unspecific activation is shown in off-target area, TH voxel (E). **(F-G)** Coronal projections featuring the main hotspots correlated with TIS-specific activation (TIS > active sham, F), and unspecific activation (TIS & active sham, G). Heatmaps represents general linear model results (z-scores) at the group level (n=5).

We next sought to explore whether TIS would produce discernible and consistent brain-wide responses, including any potential off-target effects, using fMRI based on BOLD signals. We developed a chronic electrode implantation setup, modifying a procedure optimized by Missey et al.^25^ to enable fMRI compatibility (see methods for details, **Fig S2**). First, we assessed whether the amplitude used for electrophysiological recordings (5µA) is sufficient to elicit a BOLD response. We conducted eight 30-second blocks of either 5 Hz TIS or active sham stimulation, mixed in a pseudorandomized order, with 60-second pauses in between (**Fig S3**). At a 5µA current amplitude, we were unable to detect any discernible BOLD activation. This can be attributed to the low spiking activity expected at such envelope amplitudes, in accordance with our fibre photometry results showing no neuronal entrainment below 375 µA per electrode pair (**Fig 2D**). Thus, we delivered TIS at a current amplitude of 700 µA, which elicited complete neuronal entrainment in our calcium activity study (**Fig 2D**) and was used to trigger a motor response in other investigations ^13^.

We correlated the BOLD signal with the stimulation blocks (TIS and active sham) and observed a robust hemodynamic response specifically associated with TIS in the frontal regions (IL/PL voxel in a single mouse; **Fig. 3C**). Other brain regions, such as the thalamus (TH voxel in a single mouse; **Fig. 3D**), showed BOLD signal correlations with both TIS and active sham conditions, indicating unspecific, off-target activation. To assess these effects at the group level, we used a general linear model to separately map TIS-specific and unspecific BOLD responses (**Fig. 3E-F**). Coronal projections revealed statistically significant TIS-specific activation in the IL/PL region, as well as in adjacent areas including the anterior cingulate cortex (ACA), secondary motor cortex (MOs), striatum (STR), midbrain reticular nucleus (MNR), and isolated clusters in the thalamus and hypothalamus (**Fig. 3E**). In contrast, the model of unspecific responses revealed significant clusters of both positive and negative BOLD signal in off-target regions, particularly in the thalamus and sensory cortical areas (**Fig. 3F**). The TIS-specific activation observed in frontal regions aligns with the field distributions predicted by our in silico model, while the modest involvement of striatal-thalamic circuits likely reflects engagement of downstream motor pathways ^46^; by contrast, non-specific activations in sensory-thalamic areas may indicate sensory perception of the stimulation.

Overall, our characterization of TIS spatial resolution reveals a significantly stronger neuronal engagement in the targeted region compared to off-target areas, as measured by both electrophysiological and fMRI approaches. Nonetheless, evidence of both specific and non-specific activation across the brain highlights the need to further improve the focality of TIS.

### Active field cancellation effectively dampens and steers TIS-induced neuromodulation

The cumulative findings from our dipole TI stimulation underscore the imperative for continued refinement in achieving enhanced focal precision. To do that, we decided to introduce an additional pair of electrodes that would produce a field 180 degrees shifted with respect to one of the stimulation fields of our original dipole configuration (antiphase field). We argue that by employing an active phase shift - a phase shift produced by the function generator - and an anatomy driven position of the cancellation pair, one may increase the focality of TI without requiring symmetrical electrode placement, and potentially minimizing the number of electrodes compared to previous multipair TI designs ^17,33–35,37^. We refer to the TIS with the addition of the cancelling field as *tripole configuration*.

To validate the concept of field-cancellation empirically, we started from the previously optimized protocol to target the IL region. We assumed that if the cancelling field can suppress TIS activation in the IL region, it will also be effective in dampening TI in off-target areas. The cancelling field was generated by introducing an additional independent current source with the same frequency as one of the stimulation fields (e.g., 2000Hz), but phase-shifted of 180 degrees compared to both stimulation fields (i.e., 2000 and 2005Hz) (**Fig 4A**). The cancelling pair was placed at a medial AP location, symmetrical in ML (front tripole stimulation, **Fig 4B**). We then measured the TIS field distribution in the IL *in silico* and *in vivo*, increasing the cancelling field amplitude step-wise until IL inactivation (**Fig 4B, Fig S4**). The cancelling amplitude values were then normalized by centering the 0 to the minimum TIS amplitude values, and termed *cancelling weights*, where a cancelling weight of -1 corresponds to no cancelling current amplitude, and +1 to the maximum cancelling amplitude. In the computational model, we observed a cancelling curve which showed the presence of two phenomena: cancellation and side induction (**Fig 4B-C**). Cancellation is defined as the decreasing activation of TIS in the IL region with increasing cancelling weight (normalized amplitude: -1 to 0), as the cancelling field destructively interferes with the TIS fields (**Fig 4B**, stages 1 and 2). *Side induction* refers to the constructive interference between the cancellation field and one of the TIS fields (normalized amplitude: 0 to 1) (**Fig 4B**, stage 3).

**Fig. 4.**
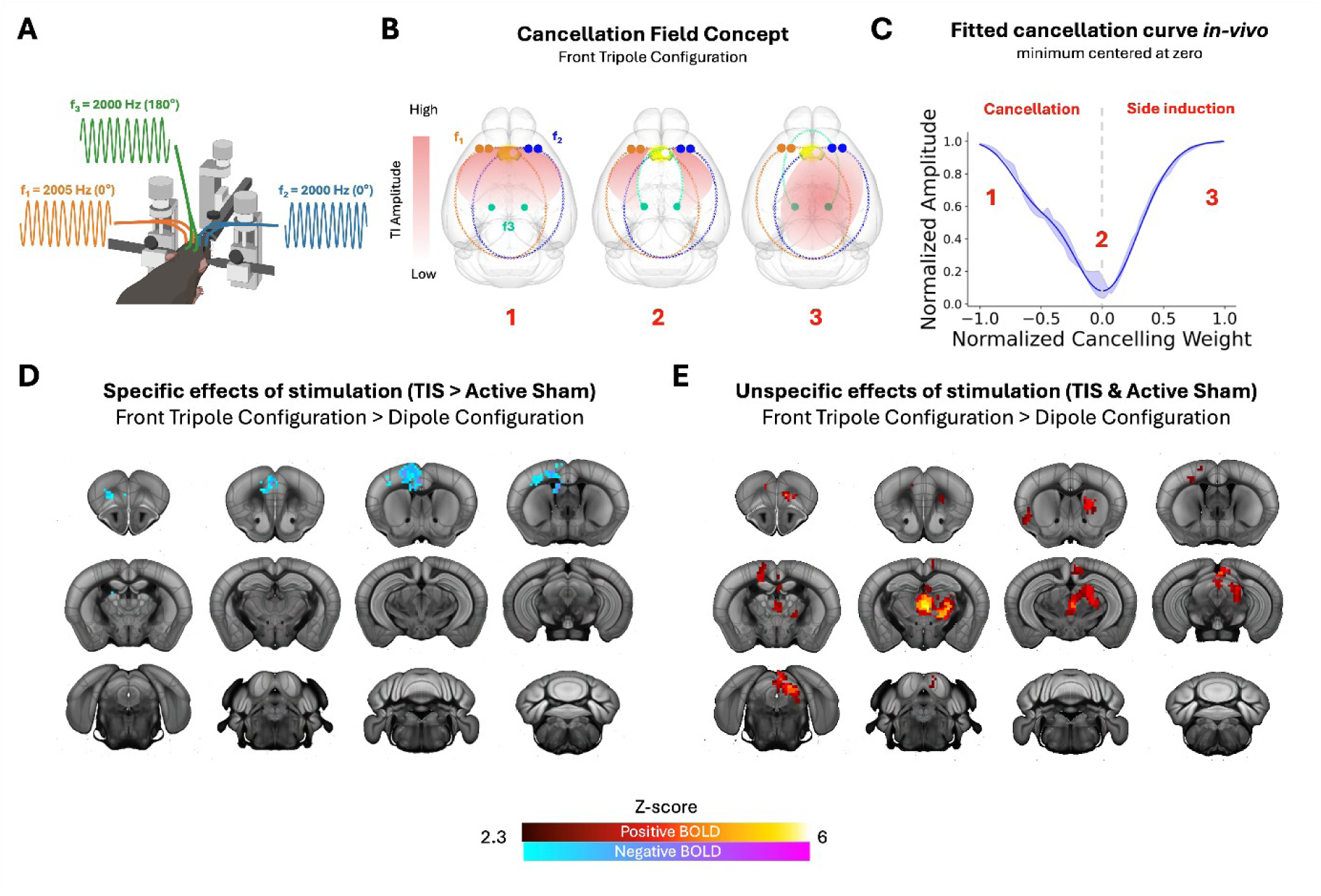
Validation of the cancelling field concept. **(A)** Schematic of tripole stimulation, with frequency andf phase for each field. **(B)** Schematic of the cancellation field concept, with illustrative TIS field distributions (in red) for each stage of the cancellation process (1-3), where electrodes and field lines are color-matched. **(C)** normalized TIS amplitude recorded in vivo in the IL region at different cancelling weights in the front tripole stimulation experiments (n=9). Solid line represents the fitted model at the group level and the shaded area depicts bootstrapped confidence interval (Q1 and Q3 percentiles). See Data Normalization section of Methods and Fig S13 for normalization details. **(D-E)** fMRI BOLD signal heatmaps extracted from a second-level analysis considering the front tripole-dipole contrast (front tripole > dipole) in the TIS-specific analysis (TIS > active sham, D) and in the unspecific signal analysis (TIS & active sham, E). Heatmaps represent general linear model results (z-scores) at the group level (n=5 for dipole and n=4 for front tripole). ACA-Anterior Cingulate Area, COL-Colliculus (includes Superior and Inferior parts), MOs-Secondary Motor Area, PL/IL- Prelimbic/Infralimbic area, RSC – Retrosplenial Cortex, STR- striatum, TH-Thalamus,

To confirm the validity of the modelling outcomes, we conducted experimental measurements *in vivo* (**Fig 4C**). We surgically implanted mice with three pairs of electrodes (two TIS pairs and one cancelling pair). During each surgical procedure, we recorded LFP activity in the IL region during TIS, to generate a cancellation curve. Consequently, for each animal, we identified the cancellation weight required to minimize TIS amplitude, corresponding to the minimum point on the cancellation curve. The resulting cancellation curve was symmetrical with respect to the minimum TIS amplitude, and well-fitted with a double Gaussian curve, at the group level (**Fig 4C**) and in individual animals **(Fig S4**). Symmetrical positioning of the cancelling pair relative to the TIS electrodes implies that each TIS field interacts with the cancelling field with a similar strength. Thus, the resulting processes of cancellation and side induction arising from these interferences are symmetrical around 0 and show comparable magnitudes of IL activation. This leads to similar contributions to the cancellation curve on both the left and right sides of its minimum, thereby producing a symmetric profile.

We reported analogous symmetrical cancellation curves with transcranial electrode positioning *ex vivo* (**Fig S5**). These curves were fitted with the same category of double Gaussian curves as the *in vivo* recordings, suggesting that the phenomenon is independent from the brain active conductance. In addition to describing the active-phase cancellation for Temporal Interference stimulation we showed that the same principle can be translated for Hz oscillations typical to tACS (**Fig S4**), allowing an additional mode of amplitude modulation for tACS (for the review of the current AM-tACS, see ^47^).

We next evaluated the effect of front tripole stimulation on brain activity using fMRI. Mice were surgically implanted with three electrode pairs to enable TI stimulation and cancellation targeting the IL region (**Fig. 4A-B, Fig. S2**). During surgery, we simultaneously recorded LFPs in the IL to generate a cancellation curve for each animal (**Fig. S6**), allowing precise determination of the cancellation weight that minimized TI-induced activity. We then applied the established stimulation and fMRI protocol to compare BOLD responses elicited by front tripole versus dipole stimulation. Using the previously described stimulation and fMRI protocol, we quantified BOLD signal during front tripole stimulation and compared it with dipole activation. We first performed an analysis of the specific TIS responses, where we mapped the brain areas with significantly different BOLD activity in the front tripole configuration compared to dipole (**Fig 4D**). We observed a significant decrease in BOLD response within the prefrontal cortical areas, especially ACC and IL, indicating an effective cancellation of brain activity in the target regions using the front tripole configuration (**Fig 4D**). Importantly, the cancellation field affected only the target area and did not eliminate TIS-induced hotspots in other regions or interfere with the effects of kHz fields. This finding further supports the spatial specificity of the cancellation approach. We then quantified the unspecific activity induced by the tripole stimulation to evaluate the potential off target effects at the whole brain level (**Fig 4E**). The tripole TIS was observed to elicit stronger unspecific BOLD changes than the dipole TIS, in particular for striatal parafasciculus and ventrolateral nuclei of the thalamus and superior colliculus. We argue that by incorporating an additional kHz field, the tripole configuration may magnify active sham-induced effects (e.g. increasing stimulus sensation) in the animals and amplifying their detection at the group level ^48^.

### Field cancellation dampens TIS off-target stimulation improving focality

After establishing the TIS cancellation principle using an anti-phase third field, we next explored its potential to suppress off-target responses and thereby enhance TIS focality. We applied the previously described TIS dipole configuration targeting the IL and added a lateral cancellation field aimed at reducing residual TIS in the ipsilateral motor cortex (MOp(ipsi), **Fig. 5A**). This region was selected as a proof of concept for testing the cancellation strategy. Since the effectiveness of this approach had already been validated using fMRI-based BOLD responses, the new electrode configuration was assessed exclusively through electrophysiology.

**Figure 5.**
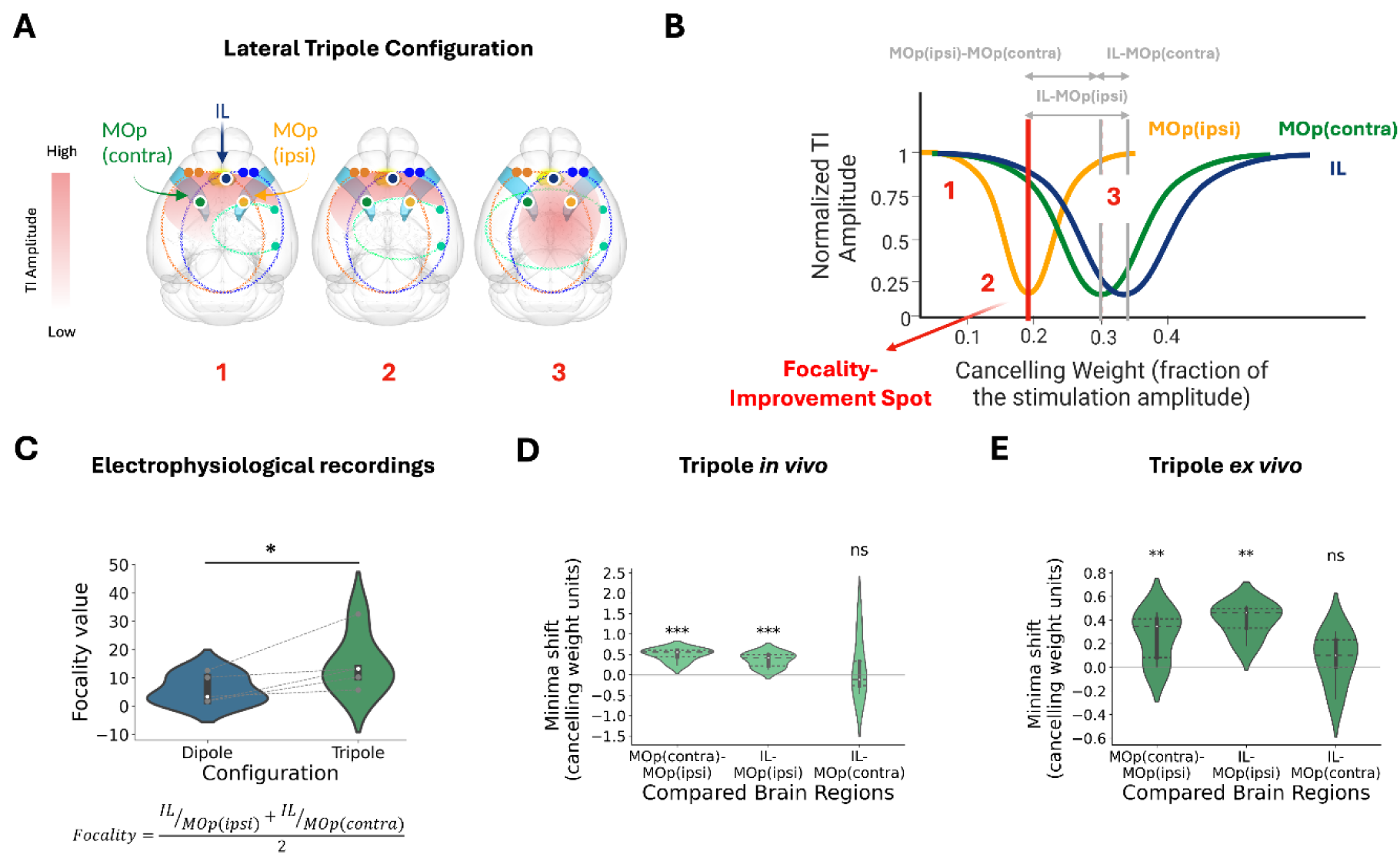
Enhancing TIS focality with phase-active cancellation. **(A)** Schematic of electrode configuration (f_1_ in orange, f_2_ in blue, f_3_ in light green) with representative TIS field distribution (in red) for each stage of the cancellation in lateral tripole stimulation experiment (1-3). The LFP recording targets are indicated as white-outlined circle (IL in blue, Mop(contra) in dark green and Mop(ipsi) in yellow). **(B)** Schematic cancellation curve for each of the recorded regions (colors match A), with a representative focality improvement spot (in red) and minima shifts indicated in grey. **(C)** Focality LFP measurements in the IL with and without cancellation field (*p<0.05, n=5, paired t-test). **(D-E)** Minima shifts between the cancellation curves from IL and MOps for the *in-vivo* (D) and *ex-vivo* (E) experiments (*p<0.05, **p<0.01, ***p<0.001. ns-nonsignificant, one-sided one sample t-tests with an alternative hypothesis of 0 being higher than a test distribution were used, p-values are FDR corrected with adaptive two-stage Benjamini-Krieger-Yekutieli method). Tested group sizes (n) are: *In-vivo*: MOp(contra)-MOp(ipsi)=6; IL-MOp(ipsi)=5; IL-MOp(contra)=5; *Ex-vivo*: MOp(contra)-MOp(ipsi)=6; IL-MOp(ipsi)=6; IL-MOp(contra)=5.

We positioned a cancellation field in proximity to the region where off-target TIS had been identified (i.e., MOp(ipsi), **Fig 3C**), creating a *lateral tripole configuration* (**Fig 5A**). To optimize the lateral tripole cancellation parameters, we placed LFP recording electrodes in three brain regions (MOp(contra), MOp(ipsi) and IL), and measured the respective cancellation curves. A cancellation curve was created by adjusting the cancellation weights to obtain the minimum TIS amplitude in the MOp(ipsi) (**Fig 5B**). Cancellation in the MOp(ipsi) required a lower cancellation weight compared to MOp(contra) and IL, because of its proximity to the cancellation electrodes, resulting in lower impedance. This method allowed us to select a focality improvement spot, where the off-target TIS activation was selectively reduced in the MOp(ipsi). The configuration obtained increased local focality in the IL region compared to the dipole configuration in *in vivo* LFP recordings (mean ± SEM: Focality(dipole)= 6±2, Focality(tripole)=15±4, **Fig 5C**). However, further increases in the cancellation weight resulted in an increase in TIS within MOp(ipsi) and reduced focality, as explained by the side induction effect.

We then measured the shifts between the cancellation curves minima with LFP *in vivo.* The MOp(contra)-MOp(ipsi) and IL-MOp(ipsi) shifts are expected to be large enough to decrease amplitude in the MOp(ipsi) without reducing it in the IL and MOp(contra) (**Fig 5B**). Conversely, IL and Mop(ipsi) minima shifts are anticipated to be small because of their similar distance to the cancelling electrode pair. We found that the MOp(contra)-MOp(ipsi) and IL-MOp(ipsi) shifts were significantly higher than zero, whereas IL-MOp(ipsi) did not show any difference (**Fig 5D**). This confirms the hypothesis that the lateral tripole configuration is a viable method to minimise TIS off-target effects without reducing the stimulation amplitude in the target.

We then replicated the experiment *ex-vivo* to decouple the effects of brain active conductance, and we were able to replicate the minima shift effect (**Fig 5E**). Interestingly, when comparing the minima shift distributions, we observed that the MOp(contra)-MOp(ipsi) distance was greater in *ex vivo* compared to *in vivo,* whereas no significant difference was observed for IL-MOp(ipsi) (**Fig S11**). This change might occur because in the *ex vivo* setup, the destruction of ion gradients in axons reduces the efficiency of signal transmission between directly connected brain areas (e.g. MOp(contra) and MOp(ipsi)), but not between areas without direct connections (e.g. IL and MOp(ipsi))^38,44^. The lower efficiency in the signal transmission led to an increase in the electrical impedance and thus, to an increase in the distance between the minima of the cancellation curves.

## Discussion

### Optimizing TIS Focality through Active Phase Cancellation

TIS has emerged in recent years as a highly promising approach for non-invasive stimulation of deep brain structures for both preclinical investigations ^13,26,49,50^ and applications in humans ^15,27,29,51,52^. However, recent studies have unveiled off-target TIS stimulation in rodents, *ex-vivo* human cadavers ^17,25,26^ and *in vivo* human participants undergoing epilepsy treatment ^41^. This poses a significant issue, as unspecific TIS could lead to unintended and confounding effects in future research and clinical applications.

In the current investigation, we adopt a multi-level approach and propose a solution to increase focality for TIS. This entailed the development of a computational modeling approach to identify the most effective electrode placements for precise targeting of the prefrontal cortex in mice with two interfering electric fields. The induced effects of dipole TIS were characterized through *in-vivo* and *ex-vivo* measurements with LFPs recordings, fibre photometry and fMRI. Together, these measurements confirmed TIS-induced neural activity and detected the presence of off-target effects at the whole-brain level, during both TIS and active sham. Such off-target effects were not eliminated via optimal parameter tuning, nor by reduction of the stimulation amplitude, highlighting the limitations of the commonly used dipole designs^15,25,26,30,50^.

We then introduced a tripolar configuration for TIS to reduce off-target effects and enhance spatial focality. This design extends the conventional dipole setup by incorporating a third electrode pair that delivers an actively phase-shifted (180°) field relative to one of the dipole components. This additional “cancellation field” selectively attenuates TI amplitudes in off-target regions while preserving stimulation in the target area. We provided a mechanistic explanation based on the spatial interplay of constructive and destructive interference and validated it through both LFP recordings and fMRI. To further confirm that the applied TIS amplitudes during fMRI were sufficient to drive neuronal activity, we performed fiber photometry, demonstrating calcium dynamics entrained to the envelope frequency. These measurements represent one of the first direct confirmations that TIS can induce neural entrainment *in vivo*, contributing critical evidence to the ongoing debate on TIS effectiveness^36,53,54^.

We report targeted spatial resolution of TIS in the IL region, where proximal off-target areas are significantly less activated than the stimulation target, in LFP recordings (**Fig 3B**, **Fig 3C**). This result is corroborated by fMRI BOLD signal (**Fig 3C**, **Fig 3E**), where a strong and specific activation is observed in the targeted area. This characterization of TIS focality constitutes a step forward in the understanding of TIS spatial resolution, showing targeted stimulation as well as other off-target cortical and subcortical areas. From our data and previous reports ^16,17,25,26,33,34,37^, it is becoming clear further efforts are required to improve focality and spatial resolution. In this study, we compared TIS-induced activation to an active sham, where both fields delivered the carrier frequency (f_1_=f_2_=2000 Hz), with the same stimulation and ramp time. This allowed us to decouple the influence of the kHz stimulation in neural activity. We observed a strong calcium response at the start of the stimulation, which decreased without affecting neural activity after 2-4 s. This onset effect of the kHz stimulation is in line with previous reports ^42^, and it is believed to depend on the ramp duration, via fast-acting membrane channel dynamics ^15^.

When examining the unspecific response—defined as brain regions activated by either TIS or active sham stimulation (**Fig. 3F**)—the BOLD signal revealed strong activation in off-target areas, suggesting that the active sham condition may exert some degree of neuromodulatory influence. These unspecific effects of TIS-induced brain activation underscore the need for researchers to exercise caution when characterizing the mechanisms of TIS. We suggest that by coupling fMRI recordings with the direct recordings of neural activity (e.g. calcium imaging^55^) one can analyze the temporal frequency of neural firing and ultimately distinguish between activations induced by TIS and effects of kHz fields. Indeed, this effect could be attributed to onset-related phenomena, which are known to be elicited by both TIS and active sham stimulation, or to alternating interference patterns of activation and conduction blocks^16^. Because the effect of the kHz frequency has not been fully elucidated, we argue for the importance of utilizing an active sham as a control in TIS studies, to detect TIS-unspecific neural activation and advance current understanding.

We propose the employment of active phase-driven TIS cancellation in already existing multipair designs ^17,33,34,37^. While such designs have advanced the focality of non-invasive stimulation by optimizing interference patterns with multiple electrode pairs, they remain limited in their capacity to directly suppress off-target activation. These methods inherently prioritize maximizing modulation at a selected region of interest. In contrast, our approach introduces a direct cancellation field, enabling selective attenuation of stimulation in specific non-targeted areas, and provides a mechanistic explanation of how this attenuation will occur. In the front cancelling field configuration, we proved that the cancelling field weakened activation in the TIS target area, using both electrophysiology and fMRI (**Fig 4**). Although the front tripole configuration is not the final application of the cancelling field concept, we argue that its ability to suppress the strongest TIS response (in the target area) can be translated to dampening weaker off-target activations. Our computational model showed that the effect of the cancelling field is largely based on direct interference between the three electric fields (**Fig 4B-C, Fig S4**). With increasing cancelling amplitude, TIS activation decreases due to negative interference until reaching a minimum point, and then it increases due to interference with one of the TIS fields (side induction). This effect is reproduced with and LFP recordings, and the cancelling is verified with fMRI, demonstrating the effectiveness and predictability of the cancelling field concept, which could serve as a tool for a range of TIS configurations. We also showed that the cancelling effect depends on the underlying brain connectome, by comparing the *in vivo* and *ex vivo* responses to TIS. This emphasizes the need to account for brain directional anisotropy in future multipair designs, to efficiently incorporate cancellation fields. The use of active, phase-driven TIS cancellation enables more flexible applications of multipair electrode designs, with the potential to reduce the number of required electric fields. Minimizing the number of applied fields is crucial due to the previously shown off-target effects associated with kHz fields ^48^, such as conduction block ^16,17,41^. Additionally, introducing an active phase-shift is a more effective solution compared to increasing the kHz frequency to reduce off-target effects ^41^. While increasing the kHz frequency may potentially decrease target TIS activation ^13,24^, as also shown in our LFP results, the cancelling field suppresses the off-target kHz fields without impacting TIS targeting.

We proposed the lateral cancellation field as an initial proof of concept for the dampening of kHz stimulation in a proximal area to the target. Our electrophysiological recordings provide evidence that the increased spatial resolution observed with the lateral tripole configuration is not due to a weaker stimulation, but rather a selective reduction of off-target stimulation. In particular, the analysis of focality demonstrates that tripole stimulation results in higher activity in the target region IL relative to the MOp off-target area (**Fig 5C**). Additionally, the shift in the minima of the cancellation curves further supports the notion of spatial selectivity, as cancellation occurs at different amplitudes in target versus off-target regions (**Fig 5D-E**). Our findings, based on both theoretical modeling and empirical evidence, establish that the incorporation of an additional phase-shifted cancellation field that can significantly enhance the spatial precision of dipole TIS. This improvement is achieved by minimizing off-target stimulatory effects, while preserving the TIS activation within the targeted cortical area. The design introduced in this study effectively addresses known limitations of TIs, opening new possibilities within the realm of its applications.

### Further prospectives for TIS-fMRI

The assessment of TIS off-target effects with LFPs relies on a-priori hypotheses driven by existing neuroanatomy and our understanding of TIS biophysics. However, the latter is currently a subject of ongoing debates and, as suggested by recent *in-silico* and *in-vivo* studies, TIS can drive changes in activity in multiple spatially distant regions ^16,17^ which can hardly be captured in their entirety with spatially restricted LFPs. Therefore, a comprehensive assessment of TIS effects should include whole-brain imaging techniques such as fMRI^56^ (for other whole-brain imaging techniques see Markicevic *et al.* ^57^). Thus, we believe that the developed MR-compatible chronic implant enabling TIS alongside fMRI scanning (i.e., TIS-fMRI) opens new avenues for TIS preclinical research. Specifically, using TIS-fMRI, we were able to show the activation of the target cortical region and associated cortico-striatal, cortico-thalamic and cortico-hypothalamic networks. Additionally, we showed that the phase-active cancellation concept can be successfully applied by using tripole TIS configuration in an fMRI setting, allowing us to assess the whole-brain effects of multipair TIS. Importantly, for the amplitude range used during fMRI experiments we additionally ensured the neuronal entrainment with fibre photometry.

An important direction that a chronic design of the MR-compatible implant unfolds is an opportunity to study TIS-induced behavioral effects and whole-brain network perturbations on the same animal and within the same experiment in case of awake rodent fMRI ^58^, mimicking the design of recent human-based TIS-fMRI experiments ^15,29^. This matching between animal and human-based experimental designs could allow the parallel application of TIS in pre-clinical and clinical studies, significantly advancing both fundamental and applied research.

The TIS setup described in this study is generalizable to rats and awake animal models. However, successful implementation requires careful consideration of several technical and behavioral factors. In awake animals, tolerability becomes critical, as suprathreshold stimulation may elicit sensory or motor responses that could confound interpretation. Moreover, adapting the setup to awake animals would require the integration of head-fixation systems and potentially revised electrode configurations to accommodate anatomical and mechanical constraints. These adaptations should be validated through complementary techniques such as modeling, electrophysiology, or optical imaging prior to deployment in fMRI. Despite these challenges, the approach holds strong potential for application in awake, behaviorally relevant settings.

In summary, our study not only empirically identified the presence of off-target TIS fields but also introduced and provided a mechanistic understanding for a novel active phase cancellation approach, enhancing TIS spatial precision. This innovation has the potential to be seamlessly integrated into current TIS protocols for both rodent and human research. Furthermore, the combination of TIS with fMRI represents a promising avenue for evaluating the effects of TIS throughout the entire brain empirically, serving as a bridge between animal-based TIS research and human studies.

### Limitations

The primary objective of this study was to empirically demonstrate a strategy for enhancing the TIS focality, supported by computational modeling. We focused on a comprehensive characterization of TIS amplitude modulation, analysing LFPs across brain regions. Whole-brain fMRI was also employed to capture global responses, and fibre photometry was used to ensure neuronal entrainment in the targeted area. An investigation by Carmona-Barron and colleagues identified c-fos and GFAP expression in blood vessel cells following TIS, introducing uncertainty about the effect of TIS on blood vessels and its potential contribution to BOLD fMRI responses ^59^.

Also, it is becoming evident in the neuromodulation field that the electrode interface plays an important role in the delivery of precise and reliable stimulation ^60^. It is known that traditional electrode materials, such as silver or platinum, are limited in their charge storage capacity, charge injection limit, and capacitive behaviour, which could create issues related to non-linear artifacts, direct current (DC) offset during stimulation, variable electrode performance and tissue damage ^61,62^. Although our electrophysiological measurements allowed us to control potential artifacts from the interface, the employment of a material with improved performances and an increased capacitive behaviour, such as PEDOT:PSS conductive polymers ^63^ or others ^60^ could reduce the risk of distorted, unreliable or off-target stimulation.

Despite these considerations, our study contributes to refining TIS techniques, offering insights applicable to humans and potential clinical applications. The TIS-fMRI setup demonstrated the whole-brain hemodynamic response to kHz fields, aligning with recent findings from computational modeling and peripheral nerve stimulation in animals ^16,17^. Furthermore, comparisons of active sham conditions between dipole and tripole designs indicated an increase in the effects of kHz fields with the number of TIS fields, potentially limiting multipair designs. Our approach, selecting the location of an additional TIS electrode based on structural anatomy and tuning the phase-shift, offers a potential strategy for optimizing multipair designs by prioritizing local focality over distant off-target stimulations in the brain. However, our results also exposed a “double-edged sword” problem, where enhancing TIS field number improves focality but may intensifies off-target effects caused by the presence of multiple kHz fields.

## Resource availability

### Lead Contact

Requests for further information and resources should be directed to and will be fulfilled by the lead contact, Valerio Zerbi.

### Materials availability

This study did not generate new unique reagents.

### Data and code availability

1. MRI data and analysis scripts reported in this paper are shared in the following Zenodo repository https://doi.org/10.5281/zenodo.15327774)
2. Any additional information required to reanalyze the data reported in this paper is available from the lead contact upon request.

## Acknowledgements

We thank Esra Neufeld and Antonino Cassara for their technical support with Sim4Life. We also thank Marius Moisa, Christopher Lewis, Friedhelm Hummel and Nicole Wenderoth for the important discussions and their support with the hardware.

## Author Contributions

Funding: VZ and RP; Design of Ephys experiments and data analysis concepts: IS, VZ, FM and AW; Design of TIS-fMRI experiments: IS, VZ and MM; Modeling of TIS fields: VB; Hardware testing: VB, IS and FM; Conduction of Ephys experiments and corresponding data analysis: IS; Fibre photometry experiments and corresponding data analysis: FC, GC, CL, CB; Conduction of TIS-fMRI experiments: IS, DK and MM; Analysis of fMRI data: IS and VZ; Computational modeling for dipole configuration with Sim4Life: VB; Computational modeling for tripole configuration with COMSOL: FM and IS; Manuscript writing: IS, SP and VZ; All authors contributed to the article and approved the submitted version.

## STAR★Methods

### Key resources table

| REAGENT or RESOURCE | SOURCE | IDENTIFIER |
| --- | --- | --- |
| Chemicals, peptides, and recombinant proteins |  |  |
| Medetomidine | Pfizer<br>Pharmaceuticals,<br>UK | Swissmedic #50590 |
| Isoflurane | Piramal Healthcare<br>Limited, India | Swissmedic #56761 |
| 1% Lidocaine | Streuli Pharma AG,<br>Schweiz | Swissmedic<br>#50564 |
| 0.9% NaCl (Saline) | B. Braun<br>Melsungen AG,<br>Germany;<br>Laboratorium G.<br>Bichsel AG, Schweiz | Item Number:<br>FE1010119 |
| Lacrinorm gel opht, eye protectant gel | Bausch&Lomb<br>Swiss AG,<br>Switzerland | Cat#7570255 |
| Lacryvisc | Alcon, Switzerland | GTIN:<br>7680499540343 |
| Fentanyl | Sintetica SA,<br>Switzerland | Swissmedic #53987 |
| Midazolam (Benzodiazepine) | Neuraxpharm<br>Switzerland AG,<br>Switzerland | Swissmedic #62556 |
| Temgesic (Buprenorphine) | Indivior Schweiz<br>AG, Switzerland | Swissmedic #41931 |
| Anexate (Flumazenil) | CPS Cito Pharma<br>Services GmbH,<br>Switzerland | Swissmedic #48280 |
| Ketoprofen | Dechra Veterinary<br>Products Suisse<br>GmbH | Swissmedic<br>#066546-01-022 |
| Bacterial and virus strains |  |  |
| AAV1-hsyn-jGCaMP8m-WPRE | Stanford University<br>vector core | Cat#169257-AAV1 |
| Deposited data |  |  |
| Allen Mouse Brain Common Coordinate Framework<br>version 3 (CCFv3) | Wang et al.<sup>39</sup> | RRID:JCR_020999<br>and<br>RRID:JRC_021000 |
| fMRI recordings | Zenodo | https://doi.org/10.5281/zenodo.15327774 |
| Experimental models: Organisms/strains |  |  |
| C57Bl/6J | Charles River<br>Laboratories,<br>Germany | C57BL/6NCrl |
| Software and algorithms |  |  |
| NiBabel toolbox | https://pypi.org/project/nibabel/ | Version 3.2.1 |
| Nilearn toolbox | https://nilearn.github.io/stable/index.html | Version 0.8.1 |
| Sklearn toolbox | https://scikit-learn.org/stable/ | Version 1.0.2 |
| SCIPY toolbox | https://github.com/scipy/scipy | Version 1.10 |
| Numpy toolbox | https://numpy.org | Version 1.20.2 |
| FSL | Analysis Group,<br>FMRIB, Oxford, UK | RRID:SCR_002823 |
| Python 3.8.13 | https://www.python.org/downloads/release/python-3813/ | N/A |
| RHX Data Acquisition Software | Intan Technologies<br>https://intantech.com/RHX_software.html | Version 3.0.4 |
| Sim4Life | Zurich Med Tech<br>AG, Switzerland<br>https://sim4life.swiss/ | N/A |
| COMSOL | Multiphysics<br>https://www.comsol.com/ | Version 5.3 |
| Doric Neuroscience Studio software | Doric Lenses<br>https://neuro.doriclenses.com/products/doric-neuroscience-studio | Version 6.1.4.0 |
| Igor Pro | WaveMetrics | Version 8.04 |

| Other |  |  |
| --- | --- | --- |
| Bruker PharmaScan 7T | Bruker Corporation, USA | N/A |
| Cryogenic coil | Bruker Corporation, USA | N/A |
| MRI-compatible small animal cradle | Bruker Corporation, USA | N/A |
| MRI-compatible heating system (LAUDA Eco E4 Silver) | LAUDA, Germany | Item number: LAU_16006 |
| Temperature probe (Optocon Fotemp4) | Optocon by Weidmann, Germany | N/A |
| MRI-compatible small animal ventilator | CWE Incorporated, USA | Part Number: 12-07000 |
| MRI-compatible small animal injectomat (Standard Infuse/Withdraw PHD ULTRA™ Syringe Pump) | Harvard Apparatus, USA | Item number: 70-3007 |
| Endotracheal intubation tubing | Multi Channel Systems MCS GmbH, Germany | Item Number: 640752 |
| Tail vein cannulation tubing | Multi Channel Systems MCS GmbH, Germany | Item Number: 640751 |
| Insulin syringe, 1 ml | Braun, USA | Omnican 100; Product Code: 9151141 |
| 2-piece fine dosage syringe, 1 ml | Braun, USA | Inject-F solo;<br>Product Code:<br>9166017V |
| Optic fibre, 0.2 mm | Doric Lenses | MFC_400/430_0.48<br>_3mm_ZF2.5(G)FLT |
| Fiber Photometry console | Doric Lenses | Item number:<br>FPC |
| LED Driver | Doric Lenses | Item number:<br>LEDD_4 |
| Fluorescence MiniCube | Doric Lenses | Item number:<br>iFMC6_IE(400-410)-<br>E1(460-490)_F1(500-540)_E2(555-570)_F2(580-680)_S |
| Mono Fiberoptic Patchcord | Doric Lenses | Item number:<br>MFP_200/230/LM<br>WMJ-0.48_1m_FCM_FCM |
| Mono Fiberoptic Patchcord - Low AutoFluorescence | Doric Lenses | Item number:<br>MFP_200/220/900-0.37_3m_FC-MF1.25_LAF |
| PVC insulated solid-core copper wire, 0.65 mm (22 AWG) diameter | Distrelect,<br>Switzerland | PRT-11367 |
| 34 AWG wires from 18-pin Wire Adapter | Intan Technologies | Part #B7600 |
| DS5 Isolated Bipolar Constant Current Stimulators | Distrelect,<br>Switzerland | N/A |
| Data acquisition card, used during Ephys | National<br>Instruments | cDAQ 9174 |
| Data acquisition card, used during Fiber photometry | National Instruments | DAQ USB-6212 |
| RHS Recording Controller | Intan Technologies | Part M4200 |
| Teflon-Coated Silver Wires 0.25 mm | World Precision Instruments, Germany | AGT1010 |
| Minipins DIL-Socket Flachprofil | RS Components, Germany | 85366930 |
| PTFE tubes 0.38 mm | Distrelec, Switzerland | TFE2-30-1220 |
| PTFE tubes 0.46 mm | Distrelec, Switzerland | TFE2-28-1220 |
| PTFE tubes 0.56 mm | Distrelec, Switzerland | TFE2-26-1220 |
| 2.5 mm heat shrinkage tube | Distrelec, Switzerland | RND 465-00234 |
| Dental cement for TI-fMRI | M+W Dental Swiss AG | Optibond Solo Plus |
| Dental cement for fibre photometry: Superbond quick monomer | Dental Addict, CH | Item number: 12-894 |
| Dental cement for fibre photometry: Superbond activateur | Dental Addict, CH | Item number: 12-810 |
| Screws for optical implant | Bilaney Consultants GmbH, Germany | Item number: P1-8L0X3905201F |
| MRI-compatible cables | neuroCare Group GmbH, Germany | N/A |
| Radio Frequency (RF) filter module | neuroCare Group GmbH, Germany | N/A |
| NeuroConn DC-Stimulator | neuroCare Group GmbH, Germany | N/A |

#### A. Experimental models and subject details

All animal experiments were performed under license from the Zürich Cantonal veterinary office following Swiss federal guidelines for the use of animals in research.

#### B. Mouse lines

For the experiments C57BL/6J mice (Charles River Laboratories, Germany) were used. Animals were housed in a group of five with 12-hour light/dark cycle and *ad libitum* food.

#### C. Methods details

##### 1. In-vivo electrophysiology

###### a) Surgical procedures

The mice were inially anesthetized with 4% isoflurane (in a 1:4 O2: air mixture as the carrier gas). The anesthesia condition was verified by assessing pedal withdrawal reflex. The mouse head was placed in a stereotaxic head frame between non-rupture ear bars on a heating pad. The animals’ body temperature (with the rectal probe) and respiration rate were monitored by the experimenter. Animals were kept in the anesthetized state with 1-2% isoflurane throughout the experiment. Prior to the start of the surgical procedure 0.1 ml of 1% lidocaine was injected subcutaneously into the scalp above the surgical region, 2-3 min later the area was shaved and disinfected with 70% Ethanol solution and the surgery started.

The scalp was opened, and the craniotomies were performed for a prospective electrode’s implantation. All coordinates were calculated using Paxinos Atlas (Third Edition, 2007). *Dura* matter was removed, and PVC insulated solid-core copper wire stimulation electrodes, (0.65 mm (22 AWG) diameter, Distrelect, Switzerland) were placed on top of the brain tissue according to the craniotomies listed in Table S2 “Front Canceling”. Recording electrodes (34 AWG wires from 18-pin Wire Adapter, Intan Technologies ©) were placed according to the stereotaxic coordinates listed in Table S2. Ground and reference electrodes were bound together and twisted before the implantation in the Cerebellum. Upon the completion of *in-vivo* recordings assigned to “Front Canceling” design, recording electrode from HIP_IPSI_ was removed and the source electrode from the cancellation pair was put at HIP_IPSI_ AP/ML coordinates on the brain surface for “Lateral Canceling” experiment (Table S2).

At the end of *in-vivo* experiments anaesthesized animals were sacrificed via cervical dislocation. After waiting for at least 1h, *ex-vivo* experiments were performed on the cadavers using the same electrode locations (Tables S2). For transcranial electrophysiological recordings electrodes from stimulation and cancellation pairs were placed on the skull surface next to the corresponding craniotomies.

##### 2. Electrical Stimulation

Three DS5 Isolated Bipolar Constant Current Stimulators (Digitimer©) were used as current sources and were driven by the input from data acquisition card NI cDAQ 9174 (National Instruments©) to produce the desired fields. For Dipole configuration one of DS5s was not used. Data acquisition card was connected to the PC allowing the manual adjustment of the field parameters via Matlab scripts.

##### 3. Electrophysiological recordings

Local field potentials (i.e. LFPs) were recorded by recording electrodes using RHS Recording Controller (Intan Technologies ©) with RHX Data Acquisition Software at 20 KHz sampling rate. The following post-processing steps were applied to the recorded time series using a custom-built Matlab script to extract TI-induced oscillations (**Fig S12**).

The principal post-processing steps included: (i) uploading the data using read_intan_RHS2000_file provided by Intan Technologies© (ii) detecting the envelope with the custom Matlab© script using a modified version of Hilbert transformation ameliorating Gibb’s distortions along with bandpass (from 0.1Hz till 250Hz) causal filtering similarly as was done in ^25^, (iii) smoothing of the detected envelope by using a sliding window of 2 sampling units, (iv) convolving the envelope with a complex Morlet wavelet at TI frequency to extract the oscillation at TI frequency ^48^,

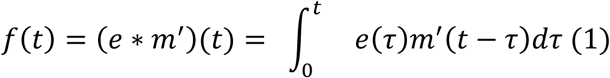

Where *e* is the extracted envelope signal, *m*′ is the Morlet wavelet and t is the analyzed recording time.

(v) f(t) analytic signal is used to extract the signal phase and amplitudes according to equations 2-3,

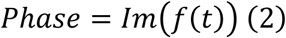

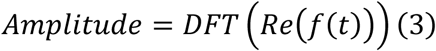

Where Im – imaginary part, Re-real part, DFT-Discrete Fourier Transform

Phase-lag index is additionally computed to verify the presence of a phase shift between oscillations from different regions. To ensure that the observed oscillation at TIS frequency (e.g. 5 Hz) is not caused by the passive intermodulation ^51^ between the high-frequency oscillations (e.g. 2000 and 2005 Hz) we verified the absence of an increased harmonic at TIS frequency in Fourier Spectrum of the raw signal (e.g. before extracting the envelope as described in step (ii)).

For the processing of tACS data, the same pipeline but without envelope extraction (step (ii)) and intermodulation detection was used.

#### D. Temporal Interference Stimulation combined with fiber photometry

##### 1. Surgical procedures

Mice were anesthetized with a mixture of isoflurane (induction 3%, maintenance 1.5%, Attane^tm^) and O_2_ during surgery and then secured in a stereotaxic frame (Stoeling). Before craniotomy, body temperature was maintained at 37 °C with a temperature controller system and Lacryvisc (Alcon, Switzerland) was applied to prevent eyes from deshydratation. For IL recording of the different neuronal subtypes (anterior posterior (AP): +1.8; medio–lateral (ML): −0.3; dorso-ventral (DV): -2; with a 10° anglemice were injected with an AAV1-hsyn-jGCaMP8m-WPRE (250 nl) produced at Stanford University vector core. During the same surgical procedure an optic fibre (0.2 mm diameter, MFC_400/430_0.48_3mm_ZF2.5(G)FLT, Doric Lenses) was implanted. Two screws were fixed into the skull to secure the optical implant, then the optic fibre was lowered 200 µm above the injection site and secure using dental cement. After surgery, mice were allowed to recover for 7 days and were habituated to handling

##### 2. Electrical stimulation

On the day of Temporal interference stimulation (TIS), mice were anesthetized with a mixture of isoflurane (induction 3%, maintenance 1.5%, Attane^tm^) and O_2_ during surgery and then secured in a stereotaxic frame (Stoeling). Before craniotomy, body temperature was maintained at 37 °C with a temperature controller system and Lacryvisc (Alcon, Switzerland) was applied to prevent eyes from deshydratation. Stimulating 0.25 teflon-coated silver electrodes (World Precision Instruments, Germany) were apposed to the dura at the desire coordinate and connected to a DS5 digitimer stimulator drive by National instrument DAQ USB-6212. At the time of stimulation, the impedance of each electrode pair was recorded. The stimulation protocol included a 500 ms ramp-up phase, followed by 10 seconds of either Active Sham or Temporal Interference Stimulation, and then a 500 ms ramp-down phase. Each trial was spaced at least 30 seconds apart and repeated a minimum of five times for each stimulation parameter (frequency or intensity).

##### 3. Fiber Photometry Data Acquisition and Processing

Fluorescence signals were recorded using a fiber photometry system (Doric Lenses Inc.) equipped with two excitation channels. A 465 nm LED (CLED_465) was used to extract Ca²⁺-dependent signals from GCaMP8m, while a 405 nm LED (CLED_405) provided Ca²⁺-independent isosbestic control measurements. Light from both LEDs was directed through a fluorescence MiniCube with 4 ports and an integrated photodetector head (iFMC4_AE(40 5)_E(460-490)_F(500-550)_S). Fluorescence emissions from GCaMP8m-expressing neurons were collected through a 200μm diameter optical fiber and directed via a detection path through a dichroic mirror to the integrated photodetector in the MiniCube.

Signal acquisition was controlled by a fiber photometry console (FPC) using Doric Neuroscience Studio software (Version 6.1.4.0). The LEDs were alternately activated in a square pulse pattern at 40 Hz, with data acquired at 12 kHz sampling rate. The acquired signals were demodulated in real-time using lock-in amplification to extract the fluorescence signals at each excitation wavelength. To calculate changes in fluorescence (ΔF/F), the 405 nm control signal was first fitted to the 465 nm GCaMP signal using linear regression to account for motion artifacts and photobleaching. The normalized change in fluorescence was then calculated as ΔF/F = (F465 - F405_fitted) / F405_fitted, where F465 represents the demodulated GCaMP signal and F405_fitted represents the fitted control signal.

The processed ΔF/F traces were exported for subsequent event-related analysis in Igor Pro (WaveMetrics), where TIS events were aligned with the fluorescence signals. For event-related analysis, we identified TIS events using an automated detection algorithm with rising-edge detection. For each detected event, fluorescence traces were extracted from a 25-second window centered around the event (-5 to +20 seconds relative to event onset).

##### 4. Spectral Analysis

The fiber photometry signal was analyzed using custom scripts in Python. For each experimental condition (Active Sham and TIS), we analyzed 5 trials per group. Time series data were collected at a sampling rate of [samples/10] Hz over a 10-second recording period. To analyze the frequency components of the signal, we performed Fast Fourier Transform (FFT) analysis. Prior to FFT computation, each trial was windowed using a Hamming window to minimize spectral leakage. The power spectrum was calculated as the squared magnitude of the FFT, normalized by the signal length, and expressed in arbitrary units. We focused our analysis on frequencies up to 7 Hz, which encompasses the analyzed frequency range. The results are presented as mean ± SEM for both the time series and power spectral density plots.

#### E. TIS combined with fMRI

##### 1. Chronic implant design

Electrodes were made from Teflon-Coated Silver Wires 0.25 mm (World Precision Instruments, Germany). One side of the wire was placed on top of the cortex, whereas the other side was used to connect to the current sources via a single minipin (RS Components, Germany) soldered to the tip of the wire. To increase the stability of the chronic implant, an additional co-axial isolation was implemented using co-axial heat shrinkage PTFE tubes (0.38 mm, 0.46 mm, 0,56mm from Distrelec, Switzerland). To avoid the lateral contact between mini-pins, each of the min-pin was isolated using 2.5 mm heat shrinkage tube (Distrelec, Switzerland). An optimal length of the wire is between 2.0 and 2.5 cm, the longer wires were more fragile and were removed by the animals during the post-surgery recovery, whereas too short wires resulted in the close position of the mini-pin to MRI cryo-coil causing distortions of the scanner magnetic field. The Teflon isolation was then removed from the wire and its tip placed on the cortex. We recommend removing the isolation not only from the wire tip, but also an additional 0.5 mm starting from the tip. This ensures a stable contact between the wire and the brain. Fig S2 A depicts an example of the used electrode with all described elements.

##### 2. Stereotaxic surgery

The mouse was initially anesthetized with 4% isoflurane (in a 1:4 O2: air mixture as the carrier gas). Intramuscular injection of a mixture of fentanyl (0.05 mg/kg), midazolam (5 mg/kg) and medetomidine (0.5 mg/kg) was performed ensuring the stable anesthesia for the whole duration of stereotaxic surgery. The following surgical procedures and craniotomies were performed as described in “*In-vivo* electrophysiology section” using coordinates from Table S2 “Front cancelling”, but with 0.25 mm Teflon-isolated silver wires (World Precision Instruments, Germany) used as stimulation electrodes. Custom-made chronic implants were fixated to the skull with dental cement (M+W Dental Swiss AG, Switzerland). Recording electrodes were connected to wires from 18-pin Wire Adapter (Intan Technologies ©) from RHS Recording Controller (Intan Technologies ©). The Teflon-isolated silver wire and the mini pin were used as recording electrodes for IL and Ground respectively (Table S2). Ground and reference electrodes from RHS Recording Controller were bound together and twisted before the connection to the minipin placed in the Cerebellum (Table S2).

LFPs recordings were performed and processed as previously discussed in “*In-vivo* electrophysiology section” section (Fig S13). We showed that the resulting TIS cancellation curve was well fitted with a double Gaussian both for individual animals and at the group level, hence reproducing the results discussed for acute electrophysiology (Fig S6). Upon the completion of the TIS cancellation experiment, the recording electrodes were removed and the remaining exposed skull surface covered with dental cement (Fig S2). Cancellation field amplitude causing the minimum TIS field in IL was used in the follow-up TIS-fMRI experiment, therefore, allowing a subject-tuned cancellation.

500 µl of Saline solution was s.c. injected to ameliorate a potential dehydration. Upon the finish of the surgery antagonist-antidote mixture was injected into the mouse consisting of temgesic (0.2 mg/kg) and annexate (0.5 mg/kg). After the surgery animals were left in a wake-up cage that has a heating pad under it until they wake up and show locomotor activity. Animals were then returned to their home cages and were single housed for the whole post-recovery phase. During the first 3 days after the surgery analgesic (Ketoprofen (10 mg/kg)) was administered daily to ameliorate a potential pain during the postsurgical recovery. At the post-surgery recovery phase all animals showed no pain symptoms, and the implant did not hamper the animal’s locomotor activity.

##### 3. TIS-fMRI experiment

###### a) MRI acquisition parameters

MRI sessions were performed using 7T Bruker BioSpec scanner equipped with a PharmaScan magnet with transmitter-receiver cryogenic coil (Bruker BioSpin AG, Fällanden, Switzerland). Adjustment procedures included wobbling, reference power and shim gradients adjustments using Paravision v6.0. Echo Planar Imaging (EPI) sequence was used for BOLD fMRI acquisition with the following parameters: repetition time (TR) = 1.5 s, echo time (TE) = 15 ms, flip angle = 60°, matrix size = 90 × 50, in-plane resolution = 0.2 × 0.2 mm2, number of slices = 20, slice thickness = 0.4 mm, and 520 volumes. Each MRI scan was accompanied by 8 Temporal Interference blocks. Each block has the following structure with either Stimulation or Active Sham condition: 60s baseline, 1s ramp-up, 30s Stimulation or Active Sham, 1s ramp-down, 1s post-baseline.

The MRI preprocessing steps, and General Linear Model (GLM) analysis were conducted using FSL FEAT (version 6) and custom-build Matlab scripts. In short, preprocessing steps included high-pass filtering (cutoff of 100s), motion correction using MCFLIRT and spatial smoothing using a Gaussian kernel with FWHM 0.5mm. The data were registered to study-specific standard space template using linear normal search and 12 degrees of freedom (DOF) options. Temporal filtering was applied on each voxel timeseries to remove the autocorrelation with FILM (FMRIB’s Improved Linear Model) (I.e. FILM prewhitening). Finally, the extracted timeseries were processed with voxel based GLM – first level analysis. The experimental design (e.g a temporal sequence of Stimulation and Active Sham conditions) convolved with double-gamma hemodynamic response function and its derivative, as well as standard motion parameters (MCFLIRT) were used as explanatory variables (EVs). All EVs underwent the same high pass filtering as was previously applied for the raw data. Z-statistics were computed with cluster thresholding p<0.05 and z >1.9. For the second-level analysis, to compare whole-brain maps between two conditions (e.g. dipole and tripole) we used fixed effects (FE) modeling feeding the results of the corresponding first-level analysis as inputs.

###### b) Hardware for Temporal Interference stimulation

Three DS5 Isolated Bipolar Constant Current Stimulators (Digitimer ©) were used as current sources and were driven by the input from data acquisition card NI cDAQ 9174 (National Instruments ©) to produce the desired fields. Stimulation electrodes from the implant were connected to the DS5 stimulators with MRI-compatible cables and electrodes (neuroCare Group GmbH, Germany) via Radio Frequency (RF) filter module (neuroCare Group GmbH, Germany). To ensure that no damage was introduced to the implant wires during the post-surgery recovery time, the impedance between stimulation pairs was measured prior to the experiment using NeuroConn DC-Stimulator (neuroCare Group GmbH, Germany). In addition, to ensure that the wires from the current stimulator and the implant were not disconnected from each other due to an accidental movement of the animal during the scan, the impedance measurements were repeated prior to every stimulation session.

###### c) Anesthesia for fMRI experiment

All mice were initially anesthetized with isoflurane (in a 1:4 O2: air mixture as the carrier gas): 4% for induction and endotracheal intubation. Endotracheal intubation was performed as previously described in ^64^. After the animal was placed in the MRI bed, it was connected to a small animal ventilator and mechanically ventilated at a rate of 90 breaths/min, with isoflurane set to 1%, with a respiration cycle of 25% inhalation, 75% exhalation, and an inspiration volume of 1.8 ml/min.

The mouse head was immobilized using an incisor tooth bar and non-rupture ear bars. Body temperature was measured using a rectal temperature probe and maintained at 36.5± 0.5°C by means of a warm-water circuit integrated into the animal holder (Bruker Biospin GmbH, Ettlingen, Germany). Following the animal preparation, the tail vein was cannulated for intravenous (i.v.) administration of a bolus of 0.05 mg/kg medetomidine (Domitor, medetomidine hydrochloride; Pfizer Pharmaceuticals, Sandwich, UK). After 10 minutes continuous infusion of medetomidine (0.1 mg/kg/h) is started. Isoflurane was lowered to 0.5%. After 20 minutes from the start of continuous medetomidine infusion MRI sessions were launched. An anesthesia solution was freshly prepared every day, by mixing saline and stock solution of medetomidine 1 mg/ml. Bolus volume and infusion rate were adjusted to the animal body weight to match the desired dose.

##### F. Computational modeling

###### 1. Dipole modeling

*<u>Model construction</u>*: While the level of segmentation improves the predictive power of the simulations, a mouse model with the highest number of segmented tissues was chosen among the models available in the Sim4Life database to find an optimal configuration of electrodes for IL stimulation. The model was generated with whole-body MR images of a 3-week male mouse (B6C3F1) and segmented into 50 elements, which included skin, fat, eye, cerebrospinal fluid, skull, muscles, tongue, and various brain structures. For the simulations, these structures were grouped into 31 tissues that had different conductivity values taken from low-frequency conductivity table of the IT’IS tissue properties database version 4.0.

The electrodes were placed on the model in accordance with the high-density EEG system used in mice, which consisted of 38 electrodes and covered the entire surface of the brain ^40^. As depicted in Fig 1A, the electrodes were represented in the simulation as cylinders with a diameter of 0.25 mm. However, it is worth noting that in the modeling pipeline, the electrodes were placed on the mouse’s scalp rather than on the *dura* matter, which is the location of the electrodes in practical experiments.

The location of the IL area in the mouse brain model was determined by co-registering the corresponding mask with Allen Brain Common Coordinate Framework (CCFv3) ^39^ which includes 213 segmented brain regions, among which bilateral infralimbic and prelimbic areas were identified as regions of interest (ROIs). Co-registration was performed in FSL 5.0 Flirt, followed by atlas alignment with the model in Sim4Life. Subsequently, surface mesh was generated from the ROIs using built-in functions in Sim4Life. The resulting highly segmented mouse model can be employed to assess the effectiveness of TI field for IL stimulation with different electrode configurations.

*<u>Quantification of TIS exposure:</u>* During TIS two distinct high-frequency waveforms, f1 (e.g., 2 kHz) and f2 (e.g., 2.005 kHz), are applied over the scalp, which differ in low frequency values – 5Hz. This results in a corresponding low-frequency amplitude modulation – hereafter referred as the TIS field or exposure, which is calculated according to equation 4:

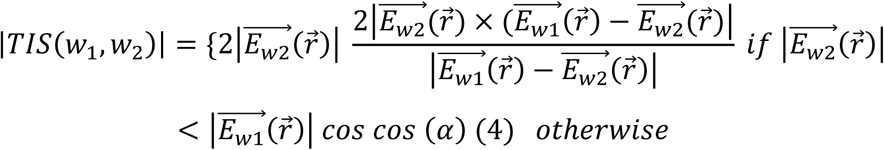

where 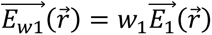, and 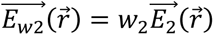. 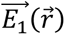 and 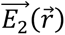 are the current normalized E-fields from both electrode pairs, while w1 and w2 are the input currents.

Identification of the most focal and efficient electrode configuration requires the calculation of E-field exposure for all electrode pairs. Instead of performing corresponding simulations for all possible pairs (N_combinations_ = 703) to save computational time, an approach based on the superposition principle was adopted that suffices with N_electrodes_-1 = 37 simulations, from which all possible exposures can be computed by linear combination. This superposition approach encapsulates the idea that one electric field generated with two electrodes can be calculated as a normalized sum of N_electrodes_-1 electric fields, where all electrodes are set as returning except for one.

Simulations of the electric fields (E-fields) were performed using ‘Electro Ohmic Quasi-Static’ finite element method (FEM) solver, which discretizes the model using adaptive, rectilinear grid. To apply the superposition principle, 37 simulations were prepared, with the Dirichlet boundary conditions being set to the electrodes. Specifically, all electrodes were assigned a stimulation of 0 V, except for one active electrode assigned a stimulation of 1 V in each simulation. The resolution of the grid was defined after a convergence analysis, that established the necessity of 0.05 mm resolution for each voxel to receive an accurate E-field calculation in the gray matter of the mouse model. Additionally, we compared the predictions of the gray matter activation in two simulations, which had different grid sizes: one of which was extended and included the head and torso of the model, while another, i.e., reduced grid, consisted only of the head of the mouse. While the predicted activations did not differ between extended and reduced models, to save the computational efforts we proceeded with generating simulations that had a reduced grid with 0.05 mm resolution, which resulted in 87 796 176 rectangular voxels per simulation. After the simulation was computed, the electric fields were down sampled to the grid with 0.1 mm resolution for each voxel to accelerate the calculation of the TIS fields.

###### 2. Tripole modeling

To model the cancellation effect of the antiphase field COMSOL 5.3 software was used. The software uses a finite element method (FEM) to solve Maxwell equations for the computation of the electrical field in the medium.

A 3D model of a mouse brain was extracted from Allen Brain Common Coordinate Framework (CCFv3). This model was used in COMSOL to simulate the mouse brain. The following parameters were used for FEM meshing: maximum element size (1.06e-4), minimum element size (7.68e-6), maximum element growth rate (1.4), curvature factor (0.4), resolution of narrow regions (0.7)

The material was chosen as water with dielectric properties adjusted to brain grey matter values at TIS carrier frequency (2000 Hz): electric conductivity (1.03e-1 S/m) and relative permittivity (9.4e+4). Stimulation electrodes were embedded as copper (COMSOL built-in properties) and placed according to the coordinates listed in the stereotaxic procedure section.

5 µA current was applied to each of the electrode pairs with a stepwise increase of the cancellation weight to produce *in-silico* results listed in Fig S5. FEM meshing as well as the scheme of the simulated field lines are shown at Fig S14.

#### G. Statistical analysis

All statistical calculations were done with custom-made Python 3.7 scripts using SciPy, NumPy, and statsmodels toolboxes. Figures were generated with custom-made Python 3.7 scripts and with BioRender.com. The exact procedures used for each analysis step are listed below.

##### 1. Data Normalization (for cancellation experiments only)

Computed amplitudes of electrophysiological recordings (see Electrophysiological recordings) from the cancellation experiments were normalized to minimize between the subject variability in absolute LFP values (see equations 5-8 and Fig S13):

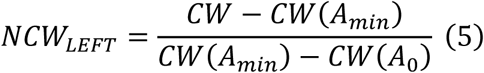

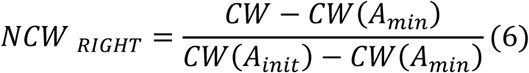

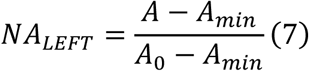

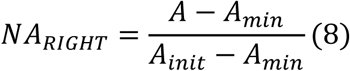

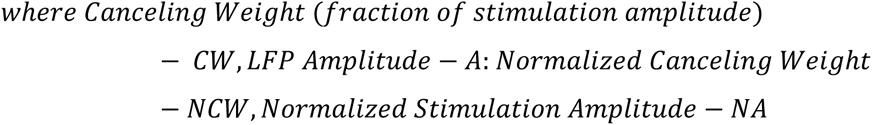

The suggested normalization brings individual U-shape curves to the common scale. Normalized Canceling weights (NCWs) vary from -1 to 1 with the minimum at 0, whereas Normalized Amplitudes (NAs) are at the range from 0 to 1. At the same time, the proposed normalization preserves the relative steepness of the cancellation curve arms (see individual data points in Fig S5 and Fig S6) which is directly linked to the studied cancellation effect.

To preserve analyzed shifts, cancelling weights were normalized only for the front tripole cancellation design.

##### 2. Group comparisons

Data distributions were tested for normality by Shapiro-Wilk test and homoscedasticity (variance homogeneity) via Levene’s test.

For n ≥3 groups, if homoscedasticity and normality of the distributions were confirmed, one-way ANOVA for independent groups or repeated measures ANOVA for paired groups were used, with unpaired or paired t-tests ^53^ for post-hoc comparisons. In case of the violation of homoscedasticity Welch corrected version of ANOVA was used. If the data were not normally distributed Kruskal-Wallis rank-sum test for unpaired or Friedman ANOVA for paired groups were used, with Mann-Whitney U-test or Quade test for post hoc comparisons.

For 2 groups, if homoscedasticity and normality of the distributions were confirmed paired or unpaired Student t-tests were used. In case of the violation of homoscedasticity, Welch version of t-test was used. If the normality condition was not met, Mann-Whitney U-test or Wilcoxon Signed Rank tests were used for comparisons between paired and unpaired groups respectively.

Two-sided tests are used by default.

False Discovery Rate was controlled by using adaptive two-stage Benjamini-Krieger-Yekutieli technique to correct p-values for multiple comparisons ^65^.

##### 3. Bootstrap

Normalized LFP data (see data normalization) from the front tripole cancellation experiments were fitted with the sum of two Gaussian functions (equation 9) at individual and group levels (e.g., pooled data).

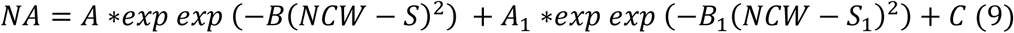

Bootstrap (N=300 iterations) confidence interval (25 to 75 percentiles) was computed to estimate the confidence of the obtained fit at the group level.

In short, for each iteration the data were resampled with the replacement keeping the size of the original sample (e.g. pooled data). Resampled data was fitted providing the values for the corresponding free parameters. The computed free parameters were used to generate the fitting curve (equation 9) with NCW step of 0.01. The NA values for the fitting curve were saved and the new iteration was launched.

##### 4. Shifts between curves minima

Formula (6) is used to calculate the shifts between curves’ minima.

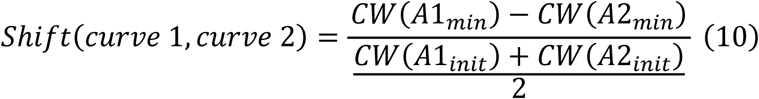

where CW(A1_min_) and CW(A2_min_) correspond to the cancelling weights at the minimum amplitudes for the curves 1 and 2 respectively and CW(A1_init_) and CW(A2_init_) are cancelling weights restored to initial values (see Fig S13 for the abbreviations corresponding to an individual curve).

Denominator at formula 10 normalizes the computed shift to the mean width of the analyzed curves.

## Supplementary Materials

**Fig. S1.**
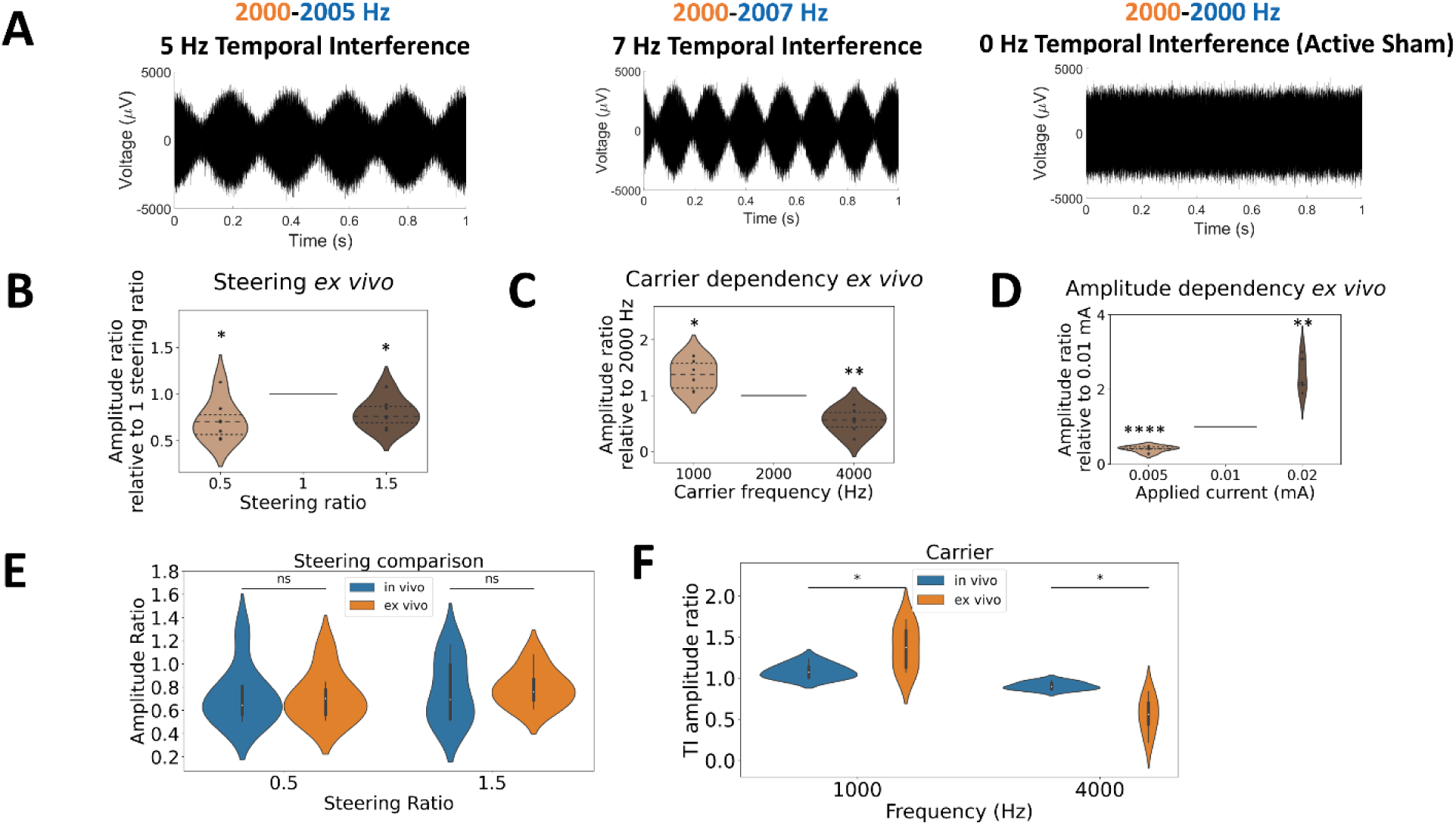
Ex-vivo electrophysiological assessment of dipole configuration. **(A)** Raw data examples of the envelope modulation (5 and 7 Hz) and Active Sham (0Hz) recorded in IL. (**B)** Steering of TI amplitude outside of IL, *p<0.05, n=7, one sample t-tests to compare amplitude ratios 0.5 and 1.5 to 1. (**C)** Dependence of TI amplitude in IL from the carrier frequency, *p<0.05, **p<0.01, n=6, one sample t-tests to compare amplitude ratios at 1000Hz and 2000Hz to 1. (**D)** Scaling of TI amplitude in IL by changing currents applied to the stimulators, ****p<0.0001, **p<0.01, n=5, one sample t-tests to compare amplitude ratios at 0.005mA and 0.02mA to 1. (**E)** Comparing TI amplitude ratios at both steering ratios, p>0.05 (ns-nonsignificant), between *in-vivo* (Fig 1E) and *ex-vivo* (Fig S1B), using U-test for 0.5 ratio and two sample t-test for 1.5 ratio (**F)** Comparing TI amplitude ratios at different carrier frequencies between *in-vivo* (Fig 1G) and *ex-vivo* (Fig S1C), for 1000Hz and 2000Hz *p<0.05, Welch’s t-tests.

**Fig. S2.**
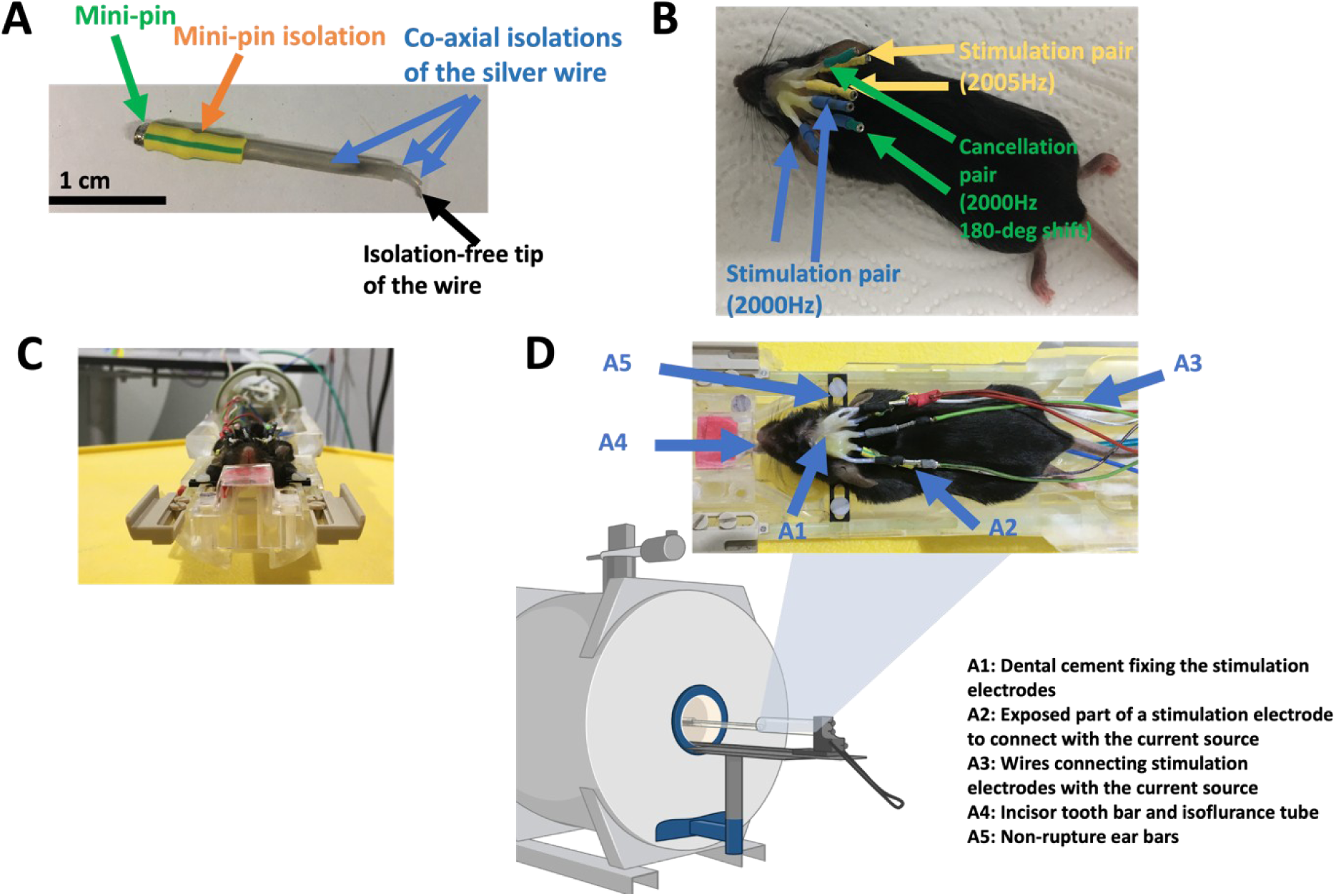
Chronic implant design. **(A)** Custom made electrodes used for the chronic implant. (**B)** Animal with the implant after the surgery. (**C-D)** Animal with the implant during the fMRI session. Different animals are shown and, therefore, colors of electrode tips are different.

**Fig. S3.**
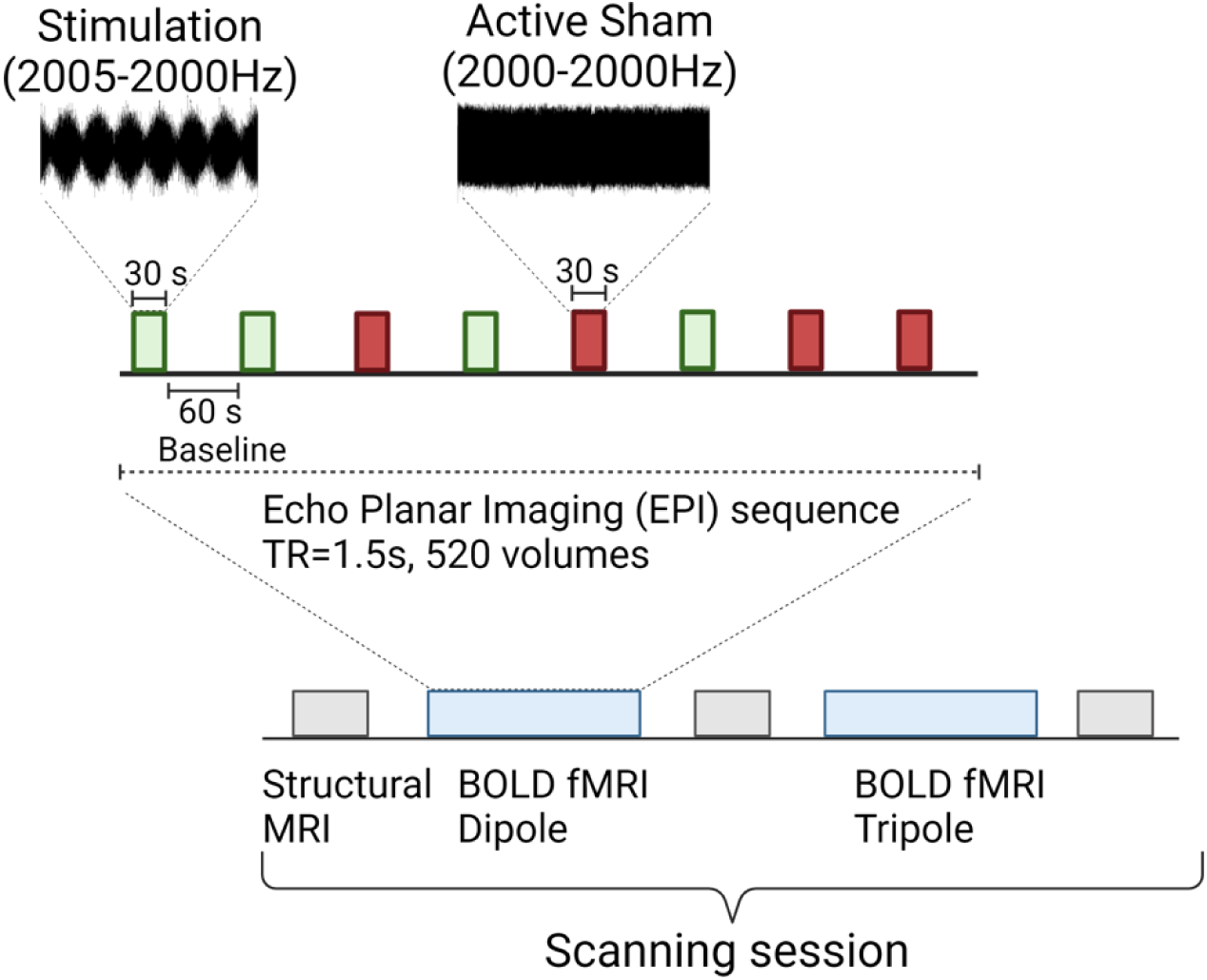
TIS-fMRI experimental procedure. TIS and active sham blocks are p0reudorandomised and delivered to each animal. The animal is scanned with an echo planar imaging (EPI) sequence, TR=1.5s, 520 volumes. The dipole and front tripole configurations are scanned on the same animal.

**Fig. S4.**
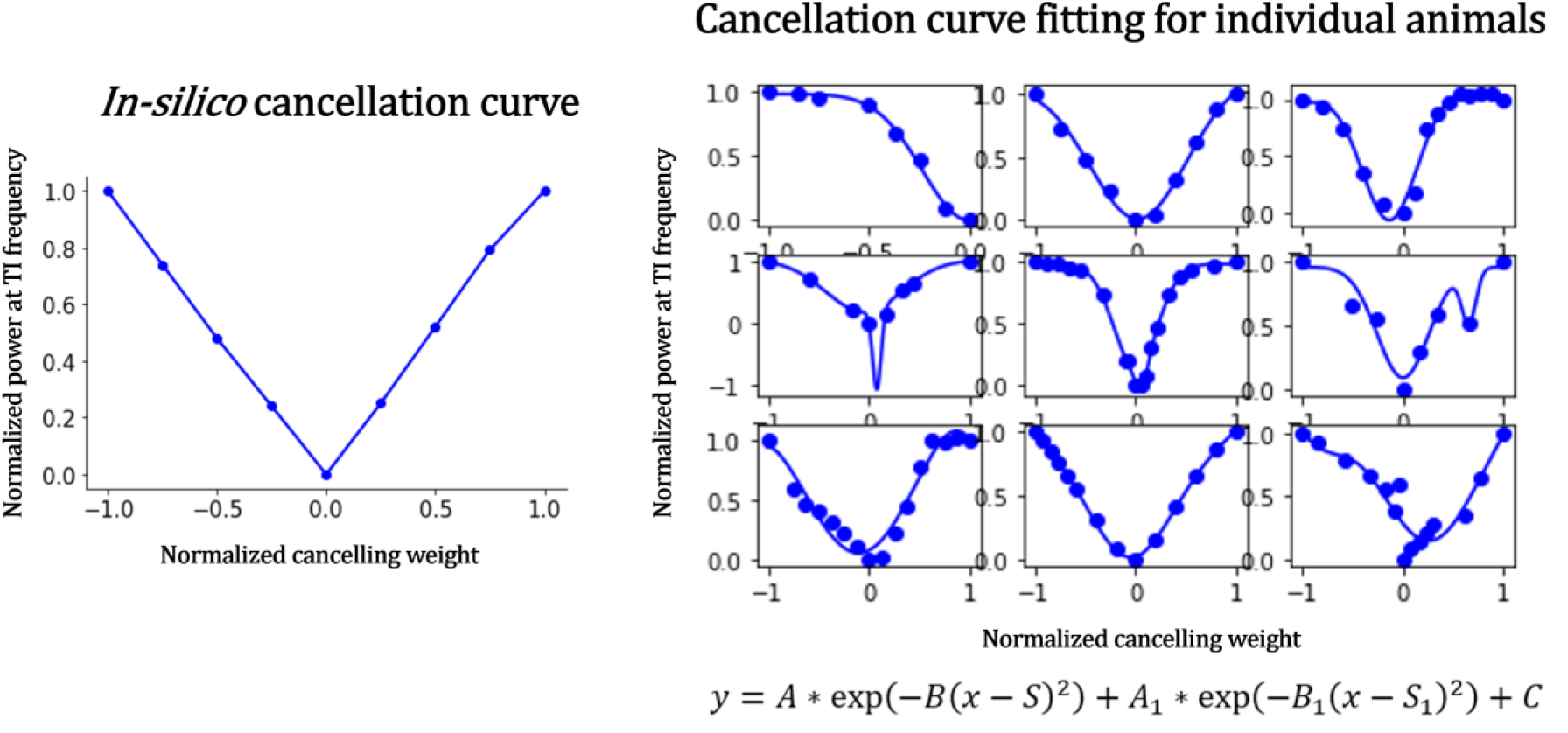
*In-silico* and individual *In-vivo* cancellation curves. Modeled cancellation curve (left) and fitted cancellation curves for individual animals *in-vivo* (n=9) (right). Each chart represents the changes in Normalized TI amplitude in IL alongside the increase of Normalized canceling weight. Dots represent individual data, whereas lines at the *in-vivo* plots (right) feature the fitting with the Double Gaussian curve (equation at the bottom).

**Fig. S5.**
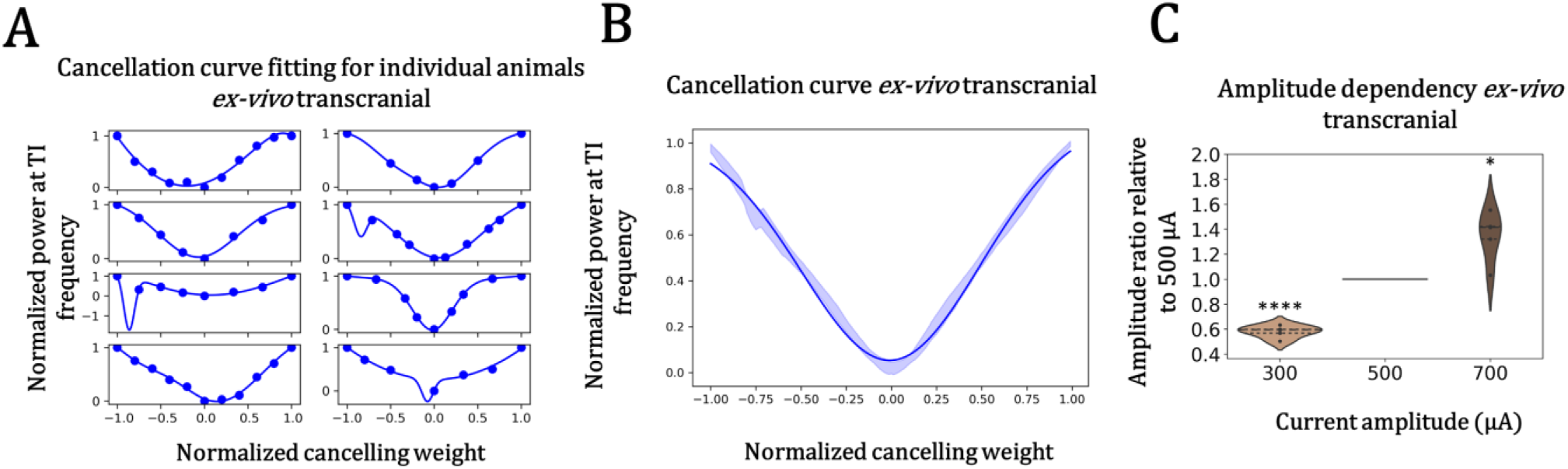
Ex-vivo transcranial TI cancellation. **(A)** Fitted TI cancellation curves (using Double Gaussian curve as in Fig. S2) for individual animals in IL (n=8). (**B)** TI amplitude in IL at different cancelling weights. Solid line represents the fitted model at the group level and shaded area depicts bootstrapped confidence interval (Q1 and Q3 percentiles). (**C)** Scaling of TI amplitude in IL by changing currents applied to the stimulators, ****p<0.0001, *p<0.05, to compare amplitude ratios at 0.3mA (n=6) and 0.7mA (n=5) to 1 one sample t-tests were used.

**Fig. S6.**
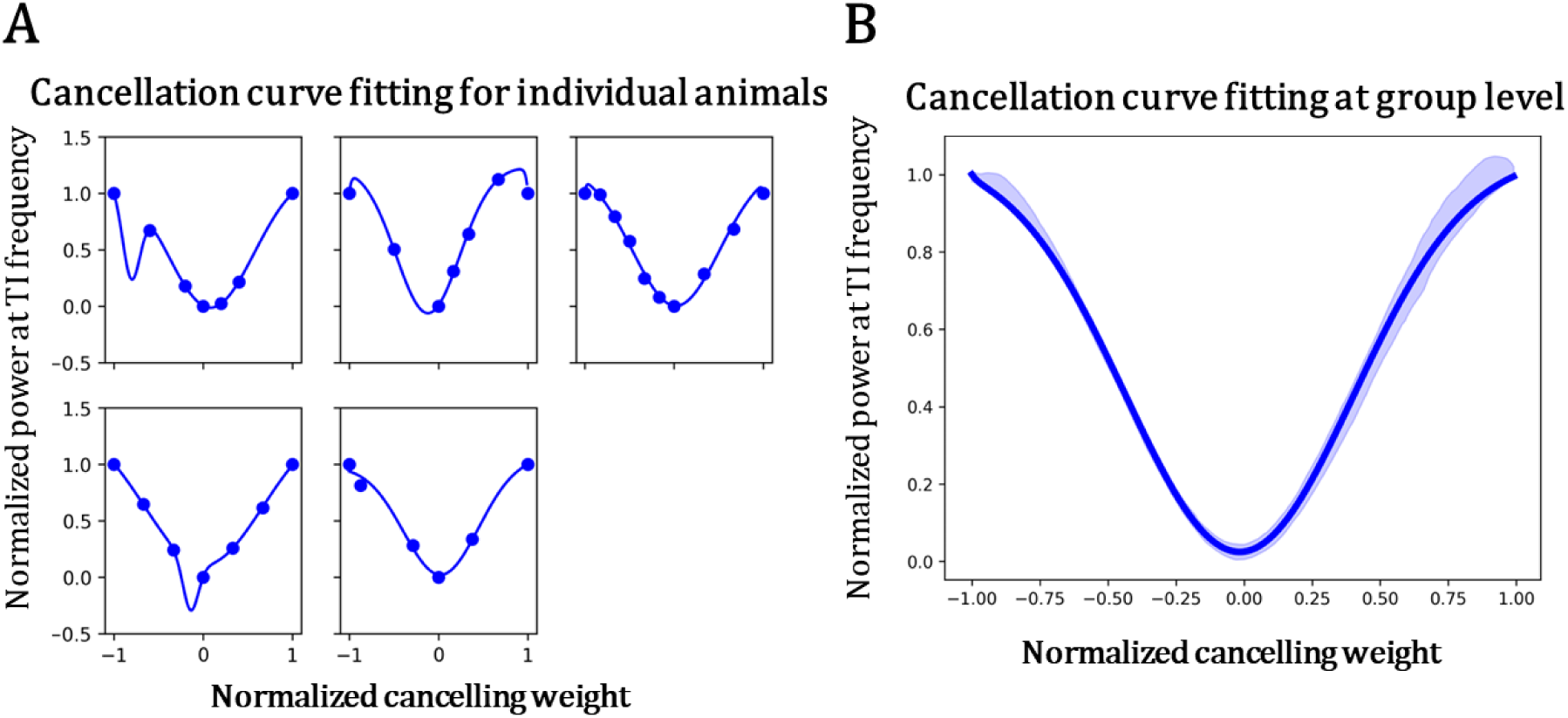
TI cancellation with a chronic implant. **(A)** Fitted cancellation curves for individual animals (n=5). (**B)** Fitted cancellation curve at group level. Solid line represents the fitted model and shaded area depicts bootstrapped confidence interval (Q1 and Q3 percentiles). Fitting was performed using Double Gaussian curve from Fig. S2.

**Fig. S7.**
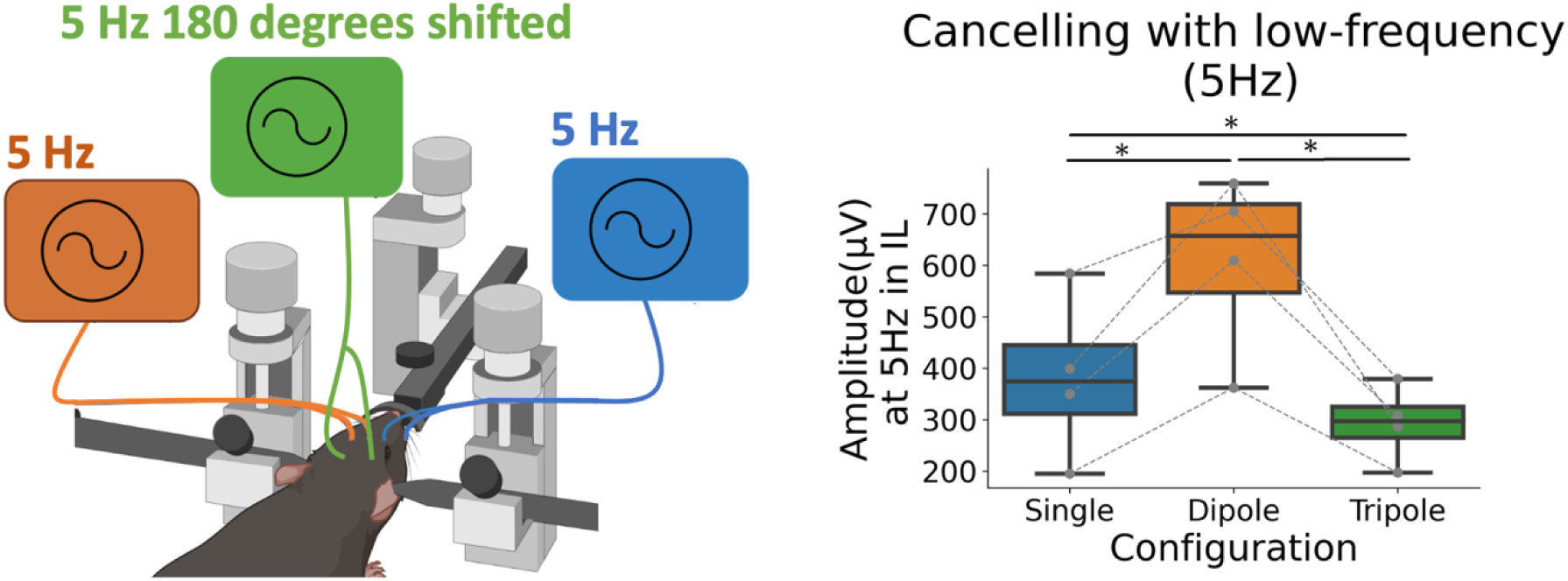
Low-Frequency cancelling field. Cancellation effect at low frequency cancelling setup. Single (Ipsilateral electrode pair), Dipole (Ipsilateral and Contralateral) or Tripole (all pairs) were used to provide 5 Hz alternating current stimulation with the same amplitude (10 µA), but with 180-degree phase shift in cancellation pair. All conditions were tested at the same animals (n=4). ANOVA followed by paired t-tests was used to check for statistical significance. p-values are corrected for FDR with with adaptive two-stage Benjamini-Krieger-Yekutieli method. *p<0.05.

**Fig. S8.**
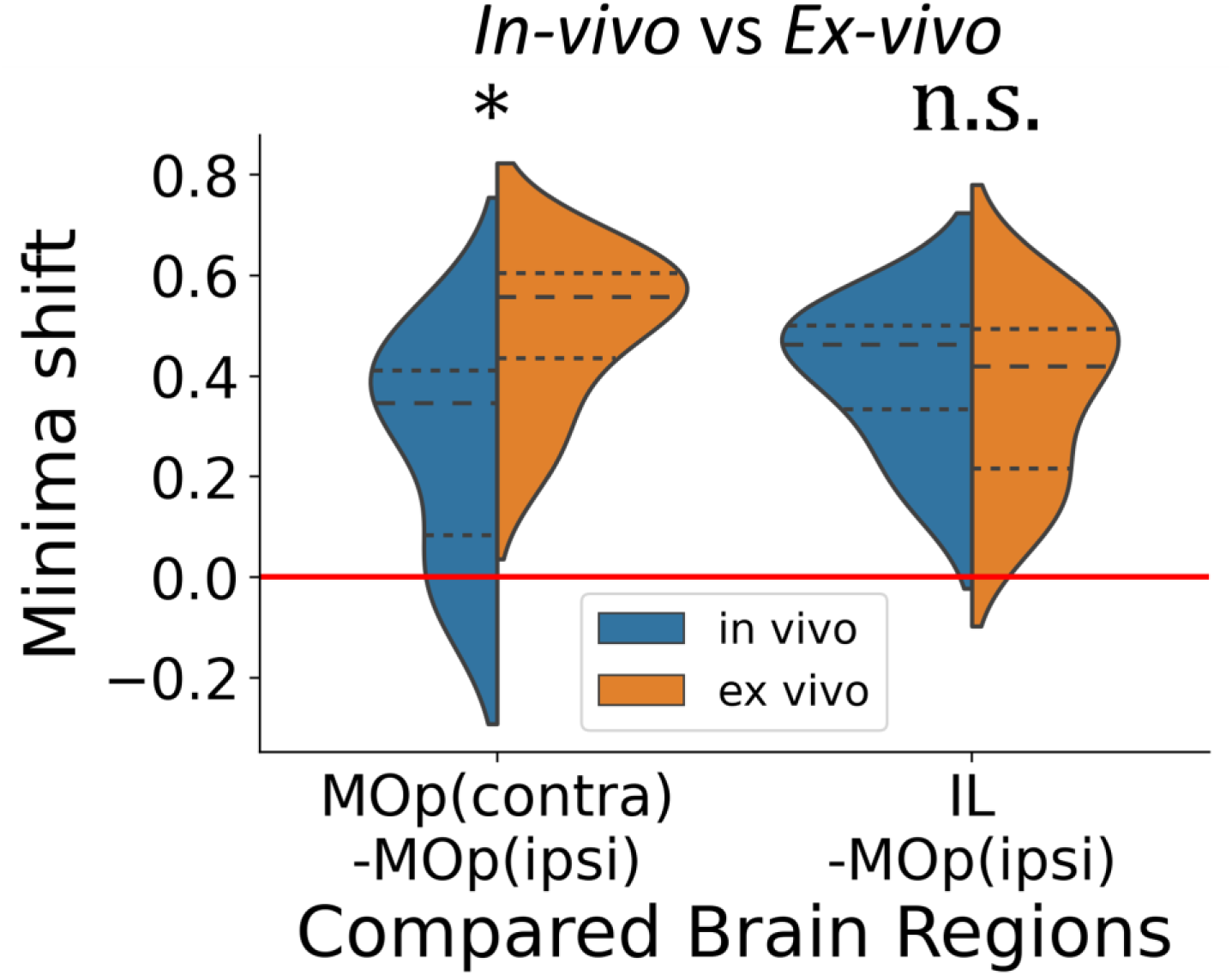
In-vivo vs Ex-vivo comparison of minima shifts. Plot outlines the comparison between minima shifts for *in-vivo* and *ex-vivo* design from Figure 2, ns-not significant, *p<0.05, t-tests. Tested group sizes (n): *In-vivo*: MOp(contra)-MOp(ipsi)=6, IL-MOp(ipsi)=5, *Ex-vivo*: MOp(contra)- MOp(ipsi)=6, IL-MOp(ipsi)=6.

**Fig. S9.**
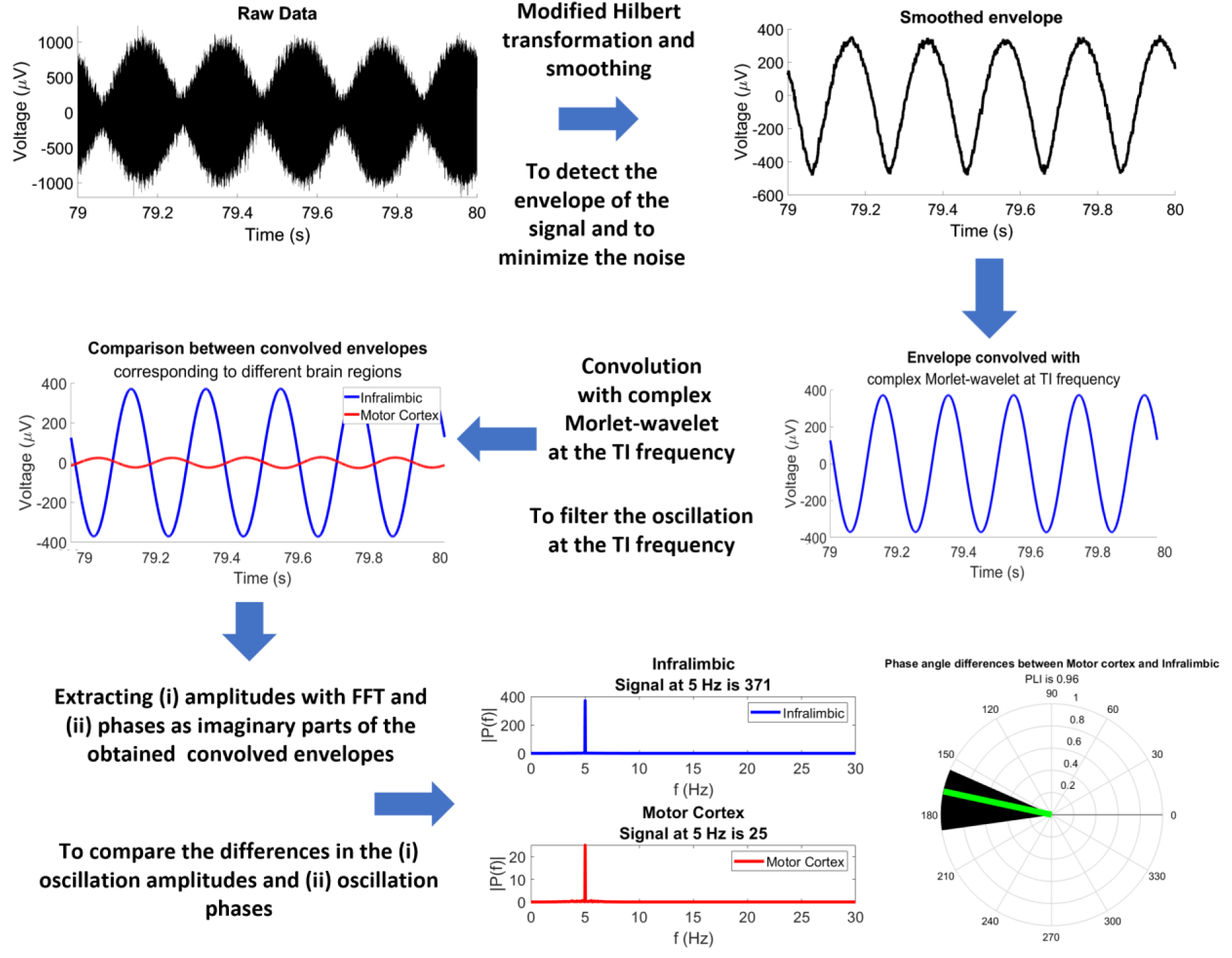
Post-processing steps of LFPs. Processing pipeline of the electrophysiological recording obtained from each individual brain areas to get the signal filtered at TI frequency (e.g., 5 Hz). Outline of the filtered signal comparison between two different brain areas (e.g. Infralimbic and Motor cortices) is also shown in the bottom panel.

**Fig. S10.**
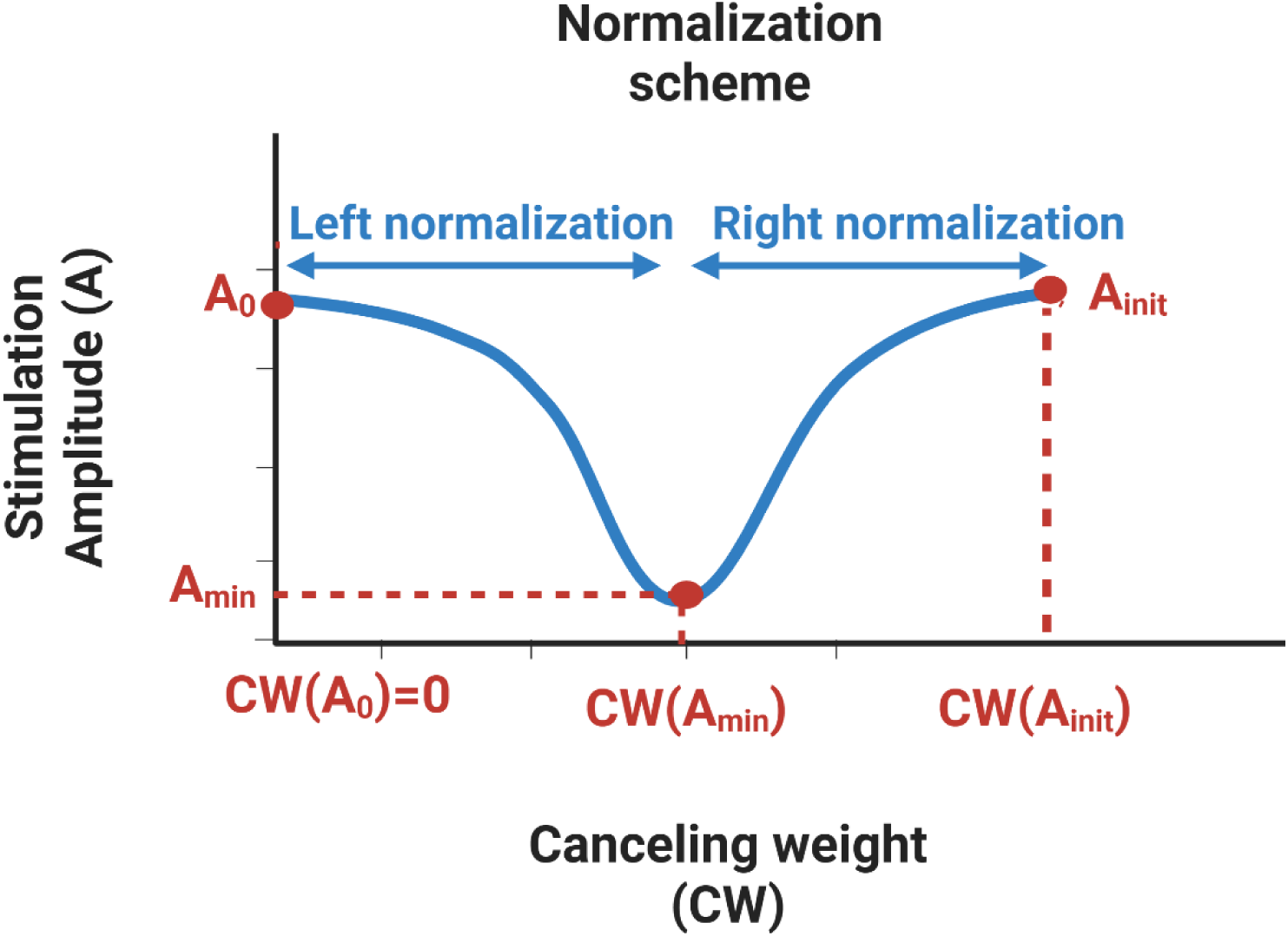
Normalization scheme. Scheme features the reference points used in the normalization of the cancellation curve at equations 5-8. A_0_, A_init_ and A_min_ refer to the Stimulation amplitudes at the corresponding canceling weights. Cancelling weight at A_0_ is 0 and corresponds to the dipole configuration.

**Fig. S11.**
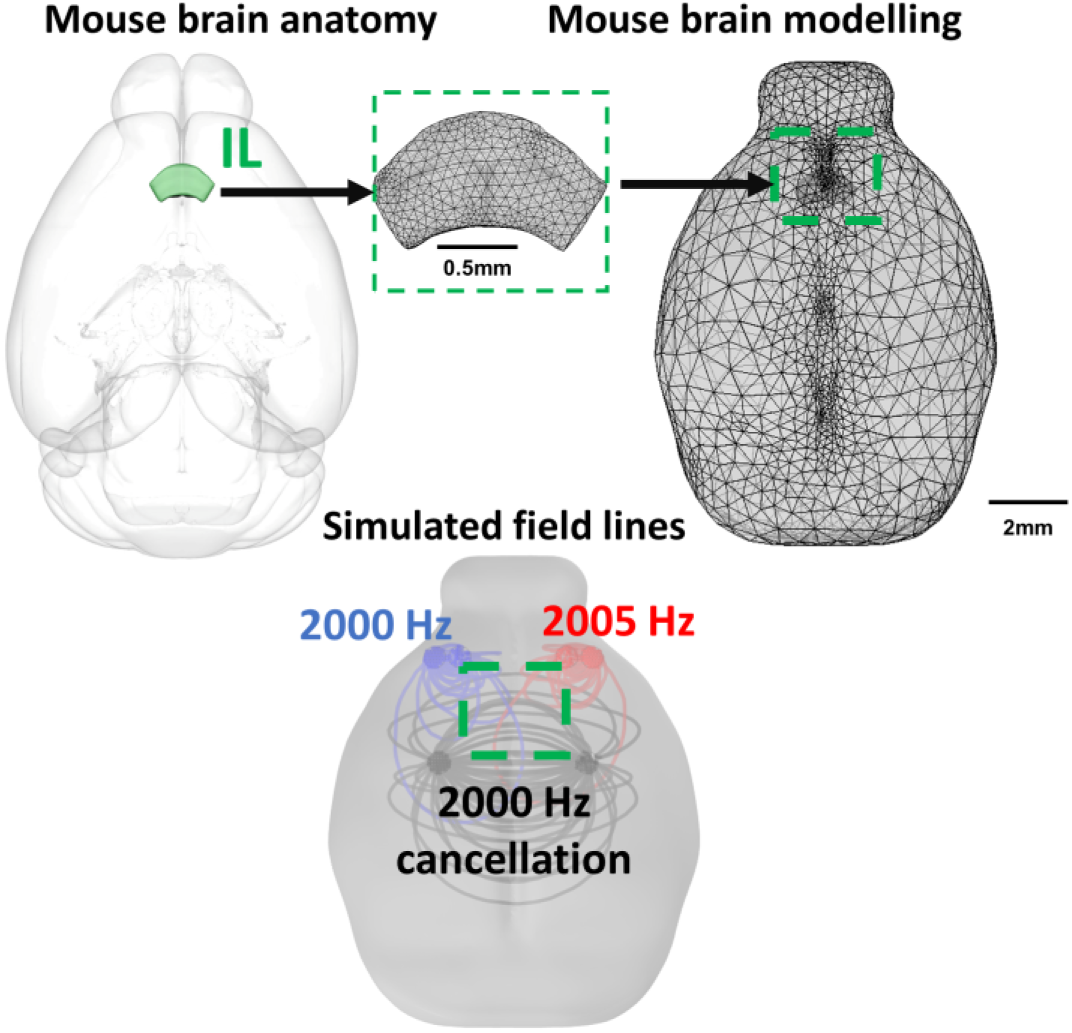
Tripole modeling scheme. Figure outlines the location of IL area in the mouse brain model and outlines FEM meshing. The bottom scheme features the modeled field lines for dipole stimulation 2000Hz(blue) and 2005Hz(red), as well as for 2000 Hz(black) cancellation field.

**Fig. S12.**
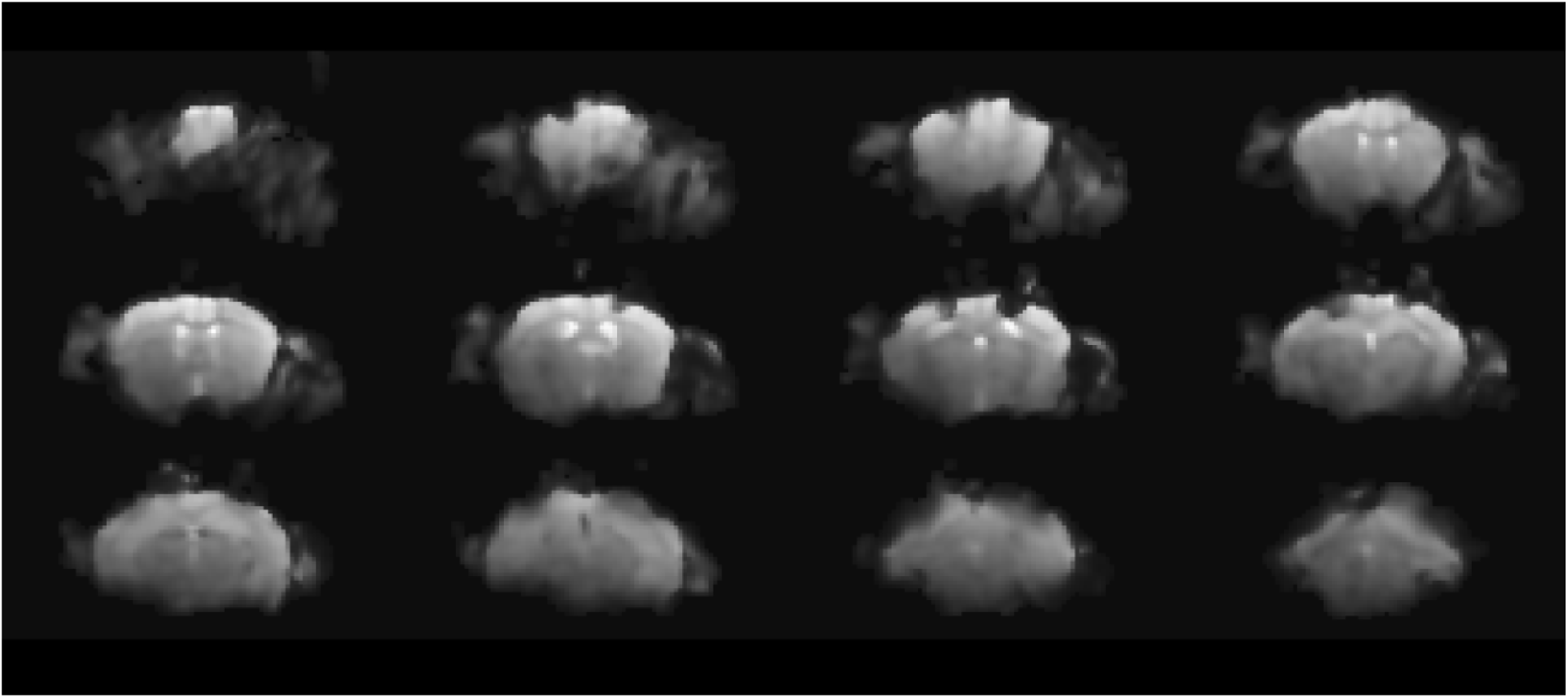
EPI-artifacts. Representative EPI images of single mouse, showing field deformations due to the electrodes

**Table S1.** TI mean phase shifts relative to IL.

| **Animals/Regions** | **MOp<sub>ipsi</sub>**<br>**(degrees)** | **MOp<sub>contra</sub>**<br>**(degrees)** | **HIP<sub>ipsi</sub>**<br>**(degrees)** | **HIP<sub>contra</sub>**<br>**(degrees)** |
| --- | --- | --- | --- | --- |
| **1** | 15 | 5 |  |  |
| **2** | 165 | 350 | 345 |  |
| **3** | 0 | 345 | 160 |  |
| **4** | 5 | 195 | 205 | 190 |
| **5** |  | 210 |  | 175 |
| **6** | 355 | 190 | 170 | 170 |
| **7** |  | 340 |  |  |
| **8** |  | 235 |  | 105 |

Each cell contains a phase shift of the TI in the corresponding region compared to IL. Empty cells correspond to the cases when the amplitude of TI was not distinguished from the background signal.

**Table S2.** TI-Ephys coordinates.

| **Electrode type** | **Electrode location** |  | **AP (mm)** | **ML (mm)** | **DV (mm)** |
| --- | --- | --- | --- | --- | --- |
| **Recording electrodes** | **IL** |  | +1.54 | -0.3 | -2.8 |
|  | **MOp<sub>IPSI</sub>** |  | -0.5 | -1 | -1 |
|  | **MOp<sub>CONTRA</sub>** |  | -0.5 | +1 | -1 |
|  | **HIP<sub>IPSI</sub>** |  | -3.1 | -2.8 | -2.5 |
|  | **HIP<sub>CONTRA</sub>\*** |  | -3.1 | +2.8 | -2.5 |
|  | **Ground (Cerebellum)** |  | -6 | -1 | -2.5 |
| **Stimulating electrodes** | **Ipsilateral Stimulation pair** | Source | +2.4 | -2.15 | surface |
|  |  | Ground | +2.4 | -1.46 | surface |
|  | **Contralateral Stimulation pair** | Source | +2.4 | +2.15 | surface |
|  |  | Ground | +2.4 | +1.46 | surface |
|  | **Cancellation Pair (front)** | Source | -0.5 | +1.75 | surface |
|  |  | Ground | -0.5 | -1.75 | surface |
|  | **Cancellation Pair (lateral)** | Source | -3.1 | -2.8 | surface |
|  |  | Ground | -0.5 | -1.75 | surface |

Stereotaxic coordinates used for the electrophysiological recordings upon TI stimulation in “Front Cancelling” and “Lateral Cancelling” designs. \* LFP in HIP<sub>CONTRA</sub> was not measured at “Lateral Cancelling design”.

